# Differential conformational expansion of Nup98-HOXA9 oncoprotein in micro- and macrophases

**DOI:** 10.1101/2025.05.21.655138

**Authors:** Hao Ruan, Rodrigo Dillenburg, Sina Wittmann, Martin Girard, Edward A. Lemke

## Abstract

Transcription factors (TFs) play a central role in gene regulation by binding to specific DNA sequences and orchestrating the transcriptional machinery. A majority of eukaryotic TFs have a block copolymer architecture, with at least one block being a folded DNA interaction domain, and another block being highly enriched in intrinsic disorder. Multivalent interactions in the intrinsically disordered region (IDR) can contribute to phase separation into macroscopic condensates. In this study, we focus on Nup98-HOXA9 (NHA9), a chimeric transcription factor implicated in leukemogenesis, in which two FG-repeat-rich IDRs derived from Nup98 get fused to the C-terminal part of transcription factor HOXA9. By integrating experiments and simulations, we examined the structural dynamics of NHA9’s FG domain across assembly states. We found that the FG domain has different conformational compactness in the monomeric state, oligomeric, and densely packed condensate state. Notably, the oligomeric state exhibits micelle-like organisation, with the DNA-binding domain exposed at the periphery. While their architecture is non-random, their sizes depend on NHA9 concentration, consistent with non-core-shell spherical micelles. Molecular dynamics simulations support the expansion behaviour of NHA9’s FG domain as oligomeric assemblies grow in size and reveal micelle-like structural features in oligomeric assemblies. These findings offer molecular insight into the phase behaviour of NHA9 and highlight the dynamic conformational transitions of IDRs during condensate formation, with implications for understanding transcriptional regulation in cancer.

## Introduction

Gene expression regulation at the level of transcription relies on the binding of transcription factors (TFs) to gene regulatory regions. TFs recognise and bind to specific DNA sequences within promoter or enhancer elements, subsequently recruiting coactivators, chromatin remodelers, and modifiers. These components collaborate to assemble the preinitiation complex which includes RNA polymerase II (RNA PoI II), thereby initiating transcription^1^. The activity of TFs is tightly regulated through various mechanisms that ultimately shape target gene expression and influence a broad range of cellular processes^2^. Eukaryotic TFs consist of multiple functional domains, among which, arguably, intrinsically disordered regions (IDRs) remain the least characterised, despite their essential roles^3^. Notably, over 80% of eukaryotic TFs contain at least one IDR, which are often substantially longer, frequently exceeding 250 amino acid residues—than those found in other eukaryotic proteins^4–6^.

Recent studies have revealed that TFs, RNA Pol II, mediator, and other transcription-associated components can self-organise into droplet-like structures frequently referred to as transcriptional condensates^7–11^. These condensates have been implicated in enhancing the spatial and temporal regulation of transcription and thus gene expression^12–19^. In many cases, multivalent intermolecular interactions mediated by IDRs within TFs can serve as a driving force for condensate formation. Intrinsically disordered proteins (IDPs, or proteins rich in intrinsically disordered regions (IDRs), which we here collectively refer to as IDPs) also promote conformational flexibility, facilitating interactions with diverse partners^8,20,21^. The abundance of IDRs is even more pronounced in chimeric transcription factors. Those are often disease-associated and formed by the fusion of the DNA-binding domain (DBD) of one gene with one or more transcriptional regulatory domains from another^22^.

In this study, we investigate the chimeric transcription factor Nup98-HOXA9 (NHA9), one of the most common Nup98 fusion proteins resulting from the t(7;11)(p15;p15) chromosomal translocation, which is associated with acute myeloid leukemia, myelodysplastic syndrome, and chronic myeloid leukemia^23–25^. Notably, ectopic expression of NHA9 has been shown to induce leukemic transformation in mouse models^26^. The N-terminal region of NHA9, derived from a component of the nuclear pore complex, contains two IDPs enriched in phenylalanine-glycine (FG) repeats, separated by a GELBS domain responsible for binding to the RAE1 protein (Fig. 1a)^27^. Recent studies have demonstrated that NHA9 undergoes phase separation to form condensates *in vitro* and hundreds of chromatin-associated puncta in the nuclei of cells^28–30^. Furthermore, NHA9 condensates has been shown to alter higher-order genome organisation, thereby modulating gene transcription during leukemogenesis^29,31^. Increasing evidence indicates that many phase-separating proteins can form heterogeneous clusters even at concentrations below the phase separation saturation threshold^32–35^. Thus, the question emerges if such nanoclusters might also exist for oncogenic fusion proteins like NHA9, how potentially diverse assembly states affect NHA9 conformation, and what functional implications this might have. While for IDPs an actual structure-function relationship as for folded proteins is not easily rationalised, the conformational ensemble is of major relevance for the IDP to execute its function. Experimentally addressing this challenge is difficult due to the large molecular size, inherent heterogeneity and flexibility of long IDPs. Similarly, tackling this computationally is challenging due to the substantial computing power and prolonged simulation times required.

**Figure 1.**
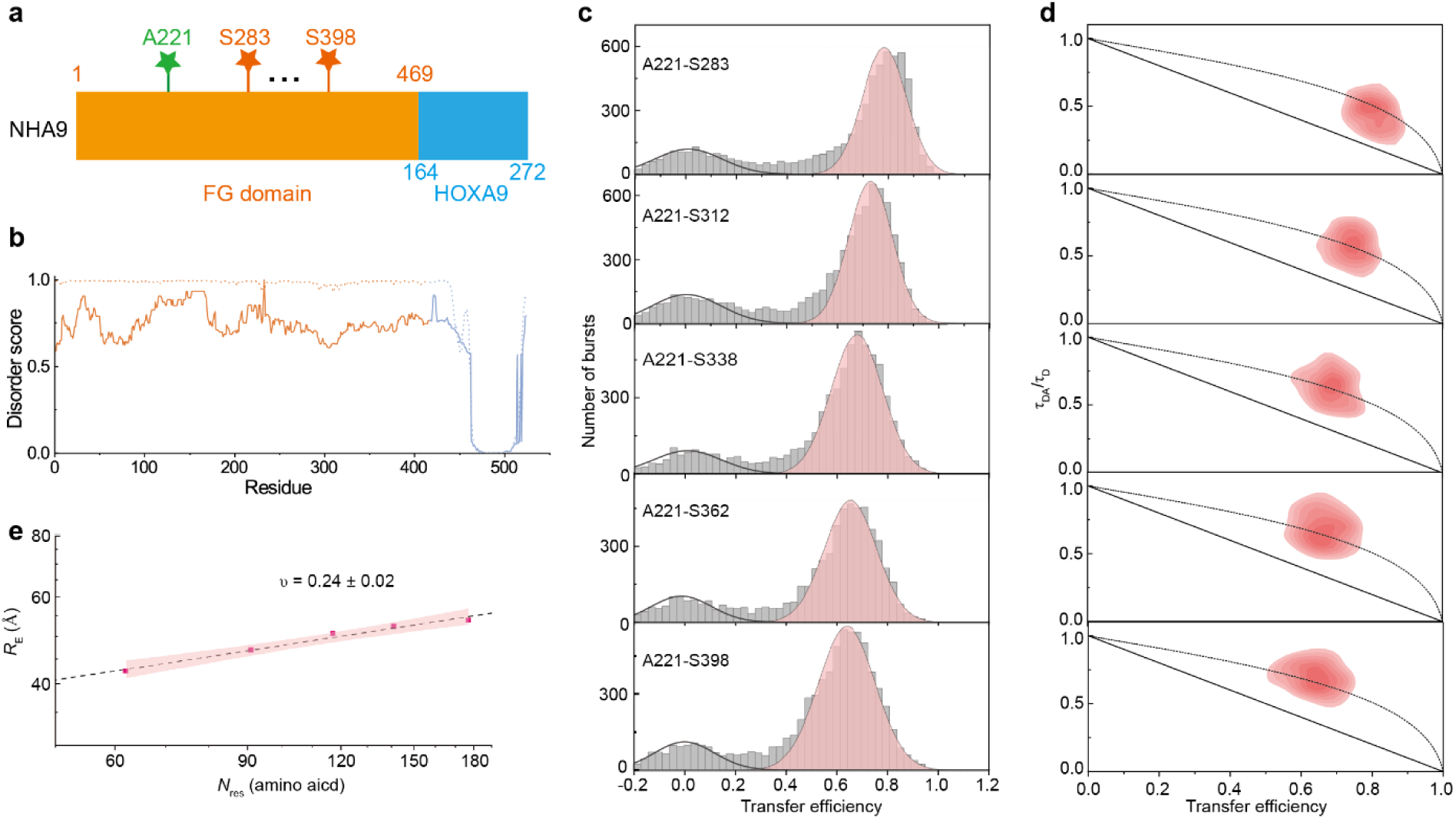
NHA9 FG domain is dynamic and highly compact in the dilute state. **a** Schematic representation of the NHA9 construct, comprising two regions: FG nucleoporins domain (yellow) and HOXA9 homeodomain (blue). Residues used for fluorescence labelling are indicated. **b** Disorder prediction as a function of residue number, calculated using two different predictors: NetSurfP-3.0^78^ (dashed line) and DEPICTER^79^ (solid line). The FG domain is highlighted in yellow and the C-terminal HOXA9 is in blue. **c** Single-molecule transfer efficiency histograms of NHA9 in the dilute state. The small peak at 〈*E*〉 ≈ 0 corresponds to donor-only labelled molecules. **d** Two-dimensional histogram of relative donor fluorescence lifetime (τ_DA_/τ_D_) versus transfer efficiency, *E*, calculated from individual bursts. The dashed line represents the dynamic line, based on a Gaussian chain polymer model. See Methods section for details. **e** Mean inter-residue distances extracted from smFRET data and scaling law fitting. The red shaded area indicates the 95% confidence interval of the fit.

Herein, we conducted a detailed characterisation of NHA9 structure and dynamics using a combination of single-molecule fluorescence resonance energy transfer (smFRET) spectroscopy, fluorescence lifetime imaging of fluorescence resonance energy transfer (FLIM-FRET), coarse-grained simulations and phase separation assays. These approaches were employed to investigate the self-association behavior of the FG domain of NHA9 across different assembly states. Our findings indicate that the FG domain of NHA9 undergoes continuous expansion while transitioning from a dilute monomeric state to an oligomeric state, and eventually to a densely packed phase. Notably, the oligomeric state of NHA9 exhibits micelle-like structural features, with the hydrophilic DBD positioned near the surface— resembling the behaviour of an amphiphilic diblock copolymer. Although their architecture is non-random and capable of binding DNA, their size depends on the concentration of NHA9, consistent with non-core-shell architectures. Taken together, the continuous transition from a dilute solution to micelle-like formation and ultimately to macrophase separation indicates that NHA9 behaves as a weak amphiphile.

## Results

### IDP of NHA9 is dynamic and highly compacted in dilute solution

Structural and disorder predictions of NHA9 suggest that the FG domains remain disordered, while the HOXA9 DNA-binding domain retains a folded and remains binding competent conformation (Fig. 1b). Structurally, NHA9 resembles a diblock copolymer, in which the disordered FG domain functions as a hydrophobic block, while the structured DBD serves as a hydrophilic block.

The disordered FG domain of NHA9 plays essential roles in activating oncogenic gene expression. To investigate this domain’s conformational behaviour and dynamics, we turned to smFRET^36–38^. A key advantage of smFRET is its ability to monitor specific subregions within structurally heterogeneous samples and to work at extremely low concentrations where aggregation phenomena are less likely to occur. We purified NHA9 from *E. coli* inclusion bodies under denaturing conditions to perform our measurements. However, the structured GLEBS domain has previously been shown to exhibit a propensity for cohesion and facilitate molecular ageing/aggregation of condensates^39,40^. Since it has been shown that deletion of the GLEBS domain does not impair NHA9-mediated transformation of primary hematopoietic stem and progenitor cells, we purified NHA9 lacking the GLEBS domain for *in vitro* experiments (Supplementary Fig. 1a)^29^. We note that even without the GLEBS domain, the protein shows a strong tendency to aggregate even at low concentrations and was thus kept in denaturing buffer until rapid dilution into physiological conditions prior to any of the following experiments. To create different FRET donor-acceptor pairs, we labelled the FG domain site-specifically by incorporating the unnatural amino acid p-acetylphenylalanine (AcF) at position 221, which was conjugated to Alexa488-hydroxylamine (donor dye), and by introducing a cysteine at various positions for labelling with Alexa594-maleimide (acceptor dye; Fig. 1a). We generated five FG domain variants of NHA9 for this analysis. The labelled proteins were analysed at the single molecule level during free diffusion through the confocal volume of the microscope to determine the mean transfer efficiency 〈*E*〉 of the FG domain. In the resulting smFRET histograms, a peak near 〈*E*〉 ≈ 0 corresponds to donor-only species, while a second population represents molecules containing functional donor-acceptor pairs. All five constructs exhibited Gaussian-shaped FRET histogram distributions with histogram widths consistent with the behaviour of a dynamic, rapidly fluctuating ensemble of IDPs (Fig. 1c).

To quantify the dimensions of the FG domain of NHA9 from smFRET data, we converted the mean transfer efficiencies into the root-mean-squared end-to-end distances (*R_E_*) between the dye pairs. This conversion was based on intrachain distance distributions modeled as a Gaussian chain, an established approximation for unfolded proteins^41,42^. To assess chain compaction independently of sequence length, we calculated the apparent Flory scaling exponent, ν, using the scaling relation *R*_E_ ∝ *N*^ν^, which relates the *R*_E_ to chain length (*N*)^43^. The FG domain of NHA9 exhibited a compact conformation in dilute solution, with a scaling exponent *ν* = 0.24 ± 0.02 (Fig. 1e), in line with the hydrophobic nature of its FG motifs leading to collapse of the protein due to strong intramolecular interactions^44^. To validate these measurements, we also performed lifetime-based analysis, which yielded *R_E_* values in line with the intensity-based analysis (Supplementary Fig. 2). To probe rapid conformational dynamics of the FG domain, we utilised relative fluorescence lifetime measurements to detect distance fluctuations between fluorophores on timescales between the fluorescence lifetime (nanoseconds) and the interphoton waiting time (microseconds). Analysis of donor fluorescence lifetimes revealed systematic deviations from the static FRET line in normalised donor lifetime versus transfer efficiency plots (Fig. 1d), consistent with a broad distribution of distances characteristic of a rapidly fluctuating IDP ensemble.

### NHA9 forms nanoclusters in subsaturated solutions with reduced compactness

Recent studies have identified that for a few IDPs, stable clusters can exist both below and above the critical concentrations at which macroscopic phase separation occurs^32,33,45–48^. To test for such an effect for NHA9, we first characterised the phase behaviour of NHA9 *in vitro* using turbidity assays. Turbidity increased sharply above around 2 μM, indicating the onset of macroscopic condensate formation (Supplementary Fig. 4a). Therefore, we focused our nanocluster measurements on concentrations below 2 μM. Due to the inherent possibility for the existence of heterogeneous nanoclusters, accurately characterising their size and conformation can be challenging. Fluorescence correlation spectroscopy (FCS) enables the determination of molecular size by analysing diffusion dynamics. We employed repetitive FCS measurements with short 10-second sampling times to reduce interference from large aggregates on the signal. Specifically, LD655-labelled NHA9 (10 nM) was used as a fluorescent probe in the presence of varying concentrations of unlabelled NHA9. Autocorrelation analysis of repeated FCS measurements yielded a distribution of average diffusion coefficients (〈*D*〉) by single-component fitting. NHA9 exhibited a bimodal size distribution, consisting of monomers (and small oligomers) and larger multimolecular nanoclusters (Fig. 2a). As the concentration of unlabelled NHA9 increased, the larger nanoclusters with heterogeneous size distributions progressively emerged.

**Figure 2.**
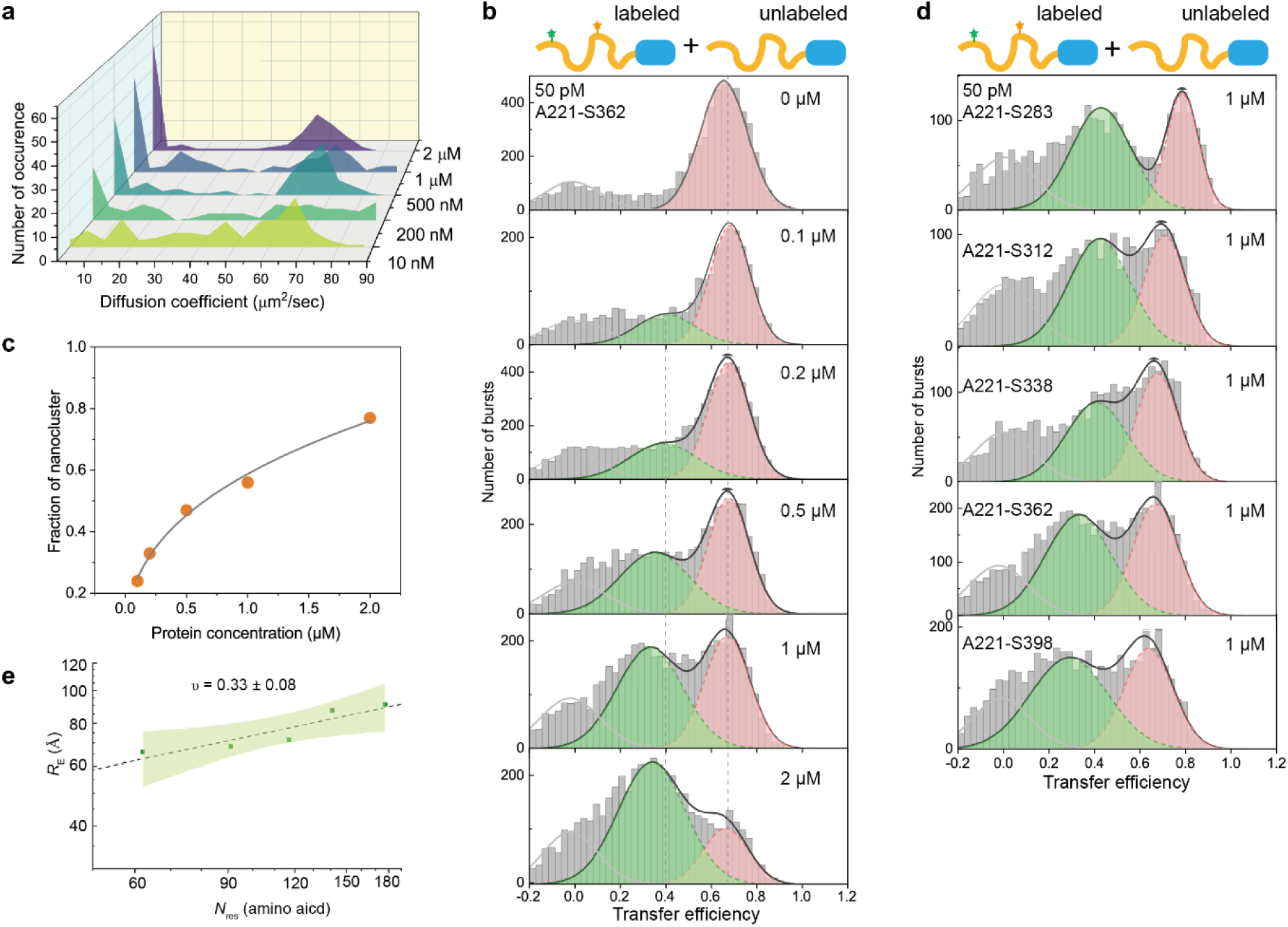
NHA9 forms nanocluster in subsaturated solutions and shows less compactness within nanocluster. **a** Concentration-dependent nanocluster formation of NHA9 measured by FCS. Autocorrelation curves were obtained from repeated (120) measurements with short sampling time (10 s), using NHA9_A221C_ labelled with LD655 as fluorescent probe in trace amounts (10 nM) with excess unlabelled NHA9. The distribution of the average diffusion coefficient at different concentrations of total protein was plotted. **b** Single-molecule transfer efficiency histograms of NHA9_A221-S362_ at increasing concentrations of unlabelled protein. The peak near 〈*E*〉 ≈ 0 corresponds to donor-only species. **c** The fraction of nanocluster extracted from Figure 2b as a function of overall protein concentration. **d** Single-molecule transfer efficiency histograms of five variants at 1 μM. **e** *R_E_* extracted from lifetime-based fitting and scaling law of the NHA9 FG domain within nanoclusters. The green shaded area indicates the 95% confidence interval of the fit.

To probe conformational changes associated with nanocluster formation, we conducted smFRET experiments using FRET-labelled NHA9 to examine transfer efficiency histograms of monomeric versus nanocluster states. Solutions containing 50 pM of FRET-labelled NHA9_A221–S362_ were doped with increasing concentrations of unlabelled NHA9. With increasing concentration of unlabelled NHA9, we observed a second peak appearing at a lower transfer efficiency, indicating the FG domain of NHA9 adopts a more expanded conformation in nanoclusters compared to monomers (Fig. 2b). The relative abundance of this low-efficiency peak increased at the expense of the higher-efficiency peak, consistent with nanocluster growth observed via FCS (Fig. 2c). Notably, the mean transfer efficiency 〈*E*〉 of the lower peak decreased from 0.40 ± 0.12 at 100 nM to 0.34 ± 0.15 at 2 μM (Supplementary Fig. 4b), suggesting a progressive reduction in compactness of the FG domain with increasing nanocluster size.

We extended these observations to four additional FG-domain variants, each with an increasing distance between the labelling sites, all of which exhibited a second, lower-efficiency peak at a concentration of 1 μM (Fig. 2d). As we also observed Acceptor quenching during nanocluster formation (see Supplementary Fig. 3), intensity-based analysis was not used to extract distance. Instead, end-to-end distances *R_E_* were obtained from lifetime-based fitting of only the donor signal using the Gaussian chain model according to Eq. 10 (see “Methods” for details). The FG domain displayed a larger apparent Flory scaling exponent (ν = 0.33 ± 0.08) in nanoclusters at a concentration of 1 μM compared to the monomeric state (Fig. 2e), indicating reduced compactness. Additionally, all five labelling variants deviated from a simple linear relationship in a 2D histogram of lifetimes versus intensity-based transfer efficiency, indicating that the FG domain also remains disordered and dynamic within the nanocluster assembly state (Supplementary Fig. 4c). In summary, combining FCS and smFRET analyses revealed a concentration-dependent increase in nanocluster size and abundance. Within the nanoclusters, the FG domain of NHA9 exhibits reduced compactness, which appears to correlate with nanocluster size.

### Impact of DNA binding on NHA9 nanocluster formation

To determine whether DNA binding affects nanocluster formation, we first confirmed the DNA-binding ability of purified NHA9 using a gel mobility shift assay (Supplementary Fig. 1b). This assay employed a 20-base-pair double-stranded DNA oligonucleotide containing the HOXA9 binding site^30^. The results confirmed that NHA9 interacts with DNA harbouring this sequence. We then investigated the effect of DNA binding on nanocluster formation using FCS. Repeated FCS measurements with short sampling intervals revealed a marked reduction in the population of larger nanoclusters upon addition of the DNA fragment (Fig. 3a). At lower protein concentration (1 μM NHA9), this effect was less pronounced (Supplementary Fig. 5), possibly due to a compensatory mechanism: the decreased number of protein molecules within nanoclusters, resulting DNA binding solubilizing NHA9 molecules from the nanoclsuter, may be offset by the added mass of the DNA fragment per nanocluster, thereby maintaining overall cluster size. In line with this, FRAP experiments also showed a slight mobility increase of NHA9 in the presence of DNA (Supplementary Fig. 5a).

**Figure 3.**
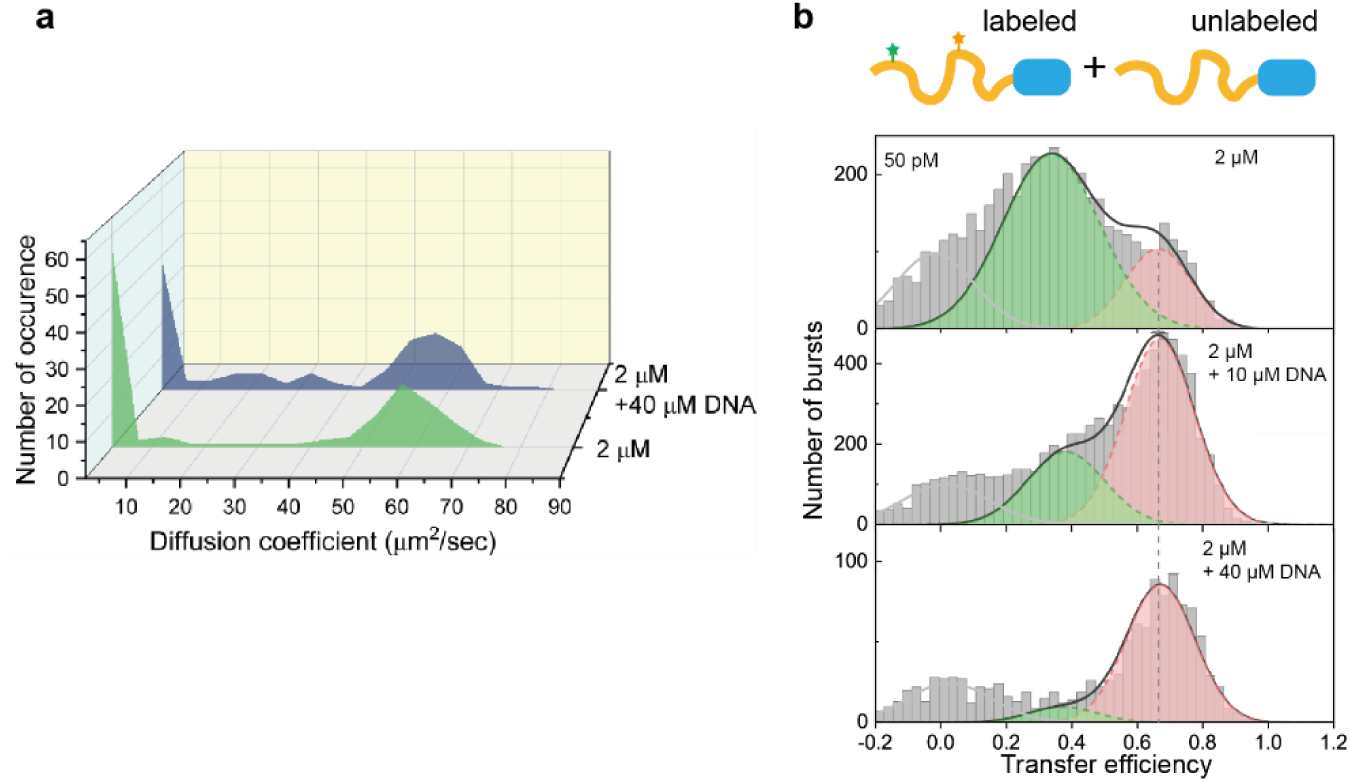
DNA influences NHA9 nanocluster formation. **a** Nanocluster formation of NHA9 at 2 µM with or without DNA fragment measured by FCS. Autocorrelation curves were obtained from repeated (120) measurements with short sampling time (10 s), using NHA9_A221C_ labelled with LD655 as fluorescent probe in trace amounts (10 nM) with excess unlabelled NHA9. **b** Single-molecule transfer efficiency histograms of NHA9_A221-S362_ at 50 pM labelled protein and 2 µM of unlabelled protein with different concentration of DNA fragment.

To further evaluate this phenomenon, we analysed smFRET histograms of NHA9 at 1 μM and 2 μM NHA9. As DNA concentration increased, the fraction of nanoclusters progressively decreased, while the monomer fraction increased, consistent with FCS observations (Fig. 3b, Supplementary Fig. 5d). Additionally, we detected a mild but consistent increase of 〈*E*〉 from 0.34 ± 0.15 to 0.37 ± 0.12 and 0.38 ± 0.1 at DNA concentrations of 10 μM and 40 μM, respectively, suggesting that DNA binding reduced the size of NHA9 nanoclusters. As a control, the 〈*E*〉 values of monomers remained constant after adding DNA (Supplementary Fig. 5c). As another control we also tested a sequence-shuffled DNA with identical nucleotide composition but lacking the HOXA9 consensus site to assess whether this effect was sequence-specific. While this control fragment also inhibited nanocluster formation, its effect was substantially weaker than that of the specific HOXA9-binding DNA as evident from the maintenance of a much larger fraction of the low 〈*E*〉 population (Supplementary Fig. 5d). These results indicate that specific interactions at least partially mediate the reduction in nanocluster size between NHA9 and its target DNA sequence. However, nonspecific nucleic acid interactions may also play a role. These findings demonstrate that DNA binding directly modulates nanocluster formation by NHA9, potentially influencing its functional activity in a cellular context.

### NHA9 undergoes significant conformational expansion in macroscopic condensates

Building on our observations of intramolecular conformational changes within nanoclusters, we investigated whether a shift in conformational distribution also occurs during phase separation. Accurate measurement of mean transfer efficiency 〈*E*〉 in protein droplets is challenging due to their inherently high background fluorescence, which can obscure the signal. To address this, we turned to fluorescence lifetime imaging of FRET (FLIM-FRET), which enables the extraction of FRET efficiency based solely on the donor’s fluorescence lifetime, thereby eliminating the need for direct acceptor excitation. Given the strong background signal in droplets, which makes FRET intensity correction more challenging, we employed ensemble FLIM-FRET measurements instead of single-molecule level measurements. The FLIM-FRET method avoids common pitfalls of intensity-based FRET methods, such as spectral spillover, and simplifies experimental design by reducing the need for extensive controls and normalisation^40,49^. To further minimise background signal, we utilised the AF594 and LD655 dye pair to label NHA9. FRET efficiency was determined by measuring the reduction in donor fluorescence lifetime upon energy transfer to the Acceptor, allowing us to estimate average spatial distances between fluorophores. Following sample preparation (see Methods), we doped a 10 μM solution of unlabelled NHA9 with 1 nM of dual-labelled NHA9 (Fig. 4a) and performed FLIM-FRET imaging of the resulting droplets.

**Figure 4.**
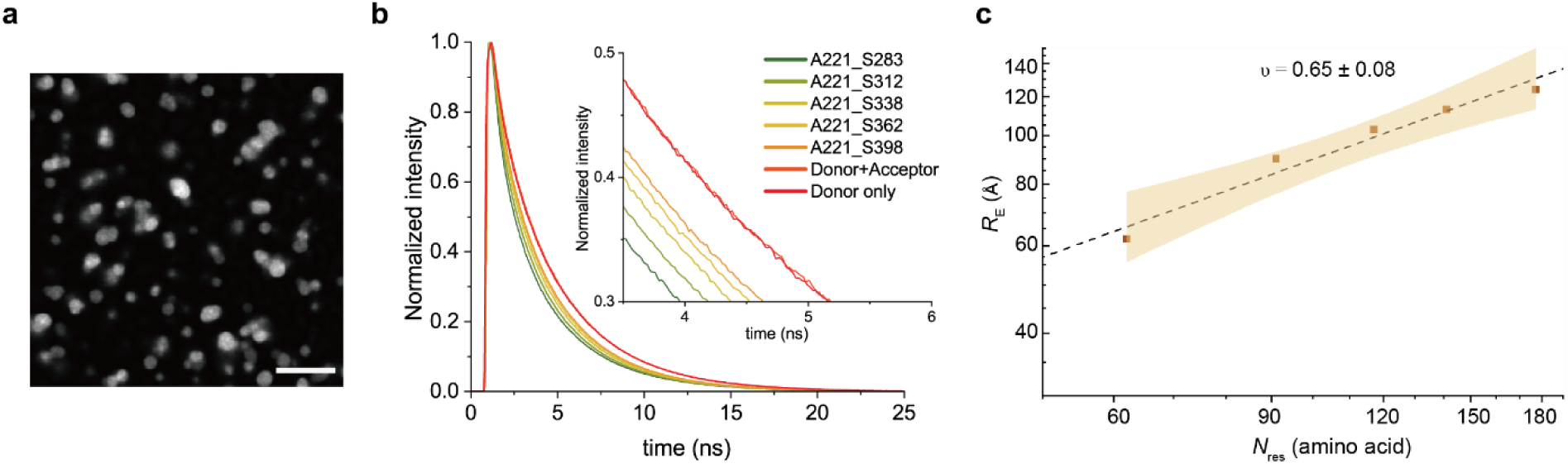
Probing dimensions of NHA9 FG domain by FLIM-FRET for *in vitro* reconstituted condensates. **a** Images of NHA9 droplets. Scale bar: 10 μm. **b** Normalised donor fluorescence decays within droplets of five NHA9 variants labelled by Alexa594-LD655 dye pair. Fluorescence decays of single mutant NHA9_A221C_ labelled with either donor only or a donor-acceptor mixture, were also included to confirm the absence of intermolecular FRET. **c** *R_E_* extracted from lifetime-based fitting and scaling law of the NHA9 FG domain within *in vitro* reconstituted condensates. The yellow shaded area indicates the 95% confidence interval of the fit.

Analysis of donor fluorescence lifetime decay curves across five NHA9 variants revealed a slower decay with increasing chain length (*N*_res_), indicating reduced FRET efficiency in longer chains (Fig. 4b). We also converted individual fluorescence decay curves into a phasor plot, which graphically represents FLIM data in a vector space. In the phasor plot, each point corresponds to the fluorescence decay profile of a single sample. Phasor analysis revealed some degree of heterogeneity among samples but also the expected overall trend, in which higher *N*_res_ exhibited a leftward shift phasor value, indicative of longer fluorescence lifetimes and increased *R*_E_ (Supplementary Fig. 6c). To rule out intermolecular FRET artifacts, control experiments were performed using single-cysteine NHA9_A221C_ mutants labelled either with the donor dye alone or with a mixture of donor and acceptor dyes. These samples were mixed with unlabelled protein at the same concentration as in the droplet assays. The fluorescence lifetime of the donor-only sample was indistinguishable from that of the donor-acceptor mixture, confirming the absence of intermolecular FRET (Fig. 4b). To quantify the *R*_E_ values, we fitted the donor fluorescence lifetime decay curves using the Gaussian chain model according to Eq. 10 (see “Methods” for details). Our analysis yielded a scaling exponent of ν = 0.65 ± 0.08 for NHA9 in the droplet phase, compared to ν = 0.33 ± 0.08 in the nanocluster state (Fig. 4c), suggesting that the FG domain undergoes significant chain expansion within phase-separated droplets. These findings indicate a progressive expansion of the NHA9 FG domain as it transitions from a dilute monomeric state to an oligomeric, and eventually, a dense condensate state.

### Probing NHA9 FG domain conformation in cells

The cellular interior presents a highly crowded and heterogeneous environment, markedly distinct from the simplified conditions typically used in *in vitro* protein studies^50,51^. Therefore, directly probing the conformational dimensions of the NHA9 FG domain within cellular condensates remains both important and technically compelling. Given that fluorophore concentrations in cells are difficult to quantify and compare between cells—and that fluorescence lifetime is independent of fluorophore concentration, acceptor emission, and instrumental fluctuations—we again employed ensemble FLIM-FRET for this study. Previously, we developed a method combining FLIM-FRET with a site-specific synthetic biology approach to probe distance distributions of disordered nucleoporins within nuclear pore complexes in live cells^40^. In this approach, FLIM-FRET is further enhanced by acceptor photobleaching, enabling precise measurement of donor-only fluorescence lifetimes and background signals.

Here, we applied this methodology to investigate the conformational dimensions of the NHA9 FG domain within transcriptional condensates in cells. Briefly, to achieve site-specific labelling, we employed genetic code expansion (GCE) to incorporate clickable non-canonical amino acids (ncAA) at defined positions (Fig. 5a). Specifically, codons in NHA9 variants were replaced with the amber stop codon (TAG) and the ncAA trans-cyclooct-2-en-l-lysine (TCO*A) was introduced to NHA9 site specifically. Fluorophores bearing tetrazine groups were then added to permeabilised cells, and labelling was achieved via strain-promoted inverse-electron-demand Diels–Alder cycloaddition between TCO*A and the dye^52^. Since TCO*A incorporation can occur in any protein containing a TAG codon, resulting in nonspecific background labelling, we utilised our recently developed orthogonally translating organelles (OTOs) (Supplementary Fig. 7). These film-like compartments accumulate both orthogonal aminoacyl-tRNA synthetases (aaRS) and the mRNA of the protein of interest (POI), thereby achieving high recruitment specificity for the NHA9 mRNA^53,54^. Furthermore, the use of small-molecule organic fluorophores for labelling ensured minimal linkage error and minimised disruption to the native structure and function of transcriptional condensates.

**Figure 5.**
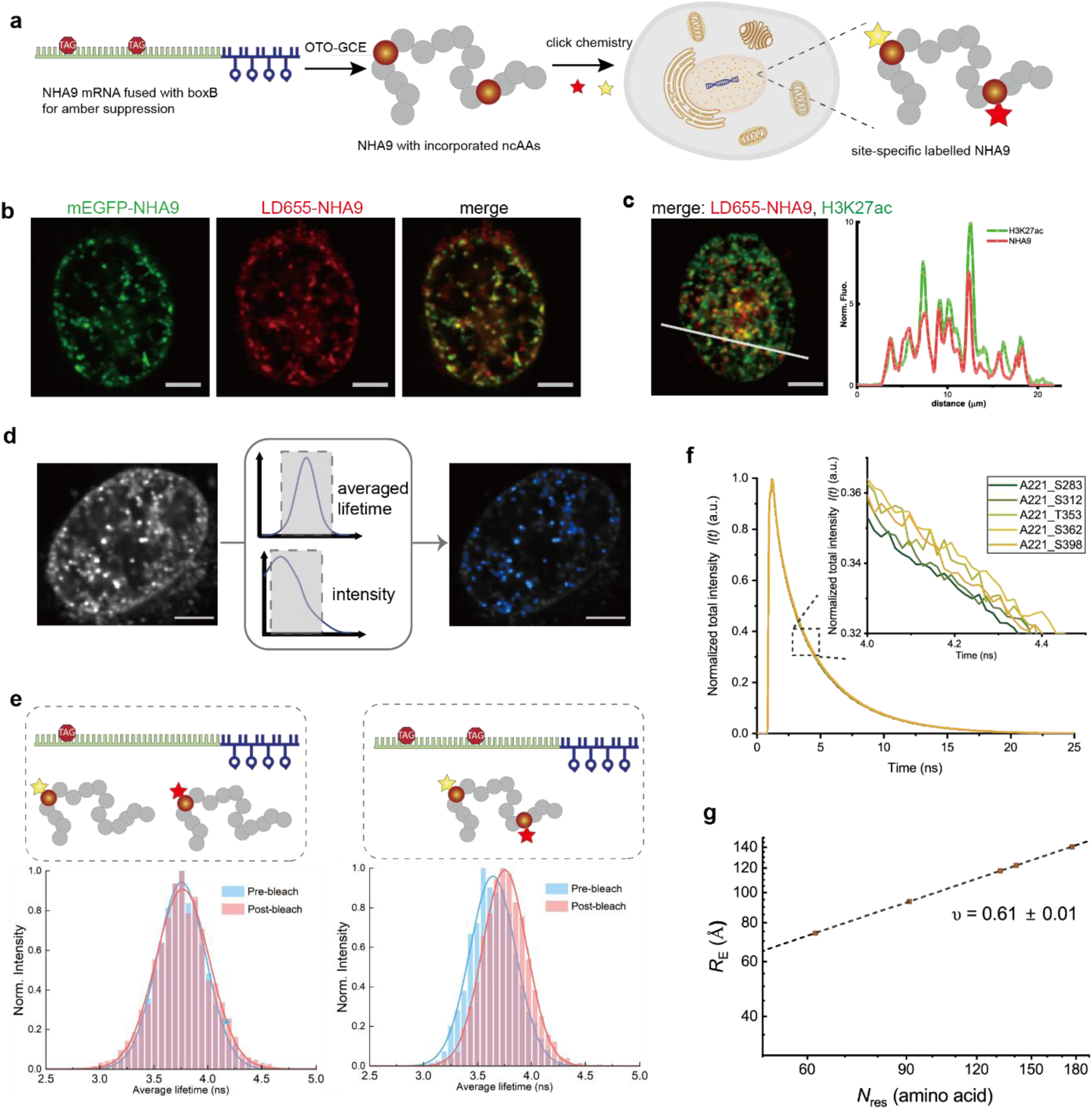
Labelling and probing dimensions of NHA9 FG domain by FLIM-FRET in cells. **a** Schematic of site-specific labelling of NHA9 in cells. NHA9 mRNA was fused with boxB loops, thus the mRNA was recruited into the OTO system. ncAAs were incorporated into NHA9 site-specifically and pair of FRET dyes was attached to NHA9 by click chemistry after adding dyes to cells. **b** Imaging for mEGFP-NHA9 fusion (green) and NHA9 labelled by LD655 via OTO-GCE (red) to assay their colocalisation. **c** Left: Imaging for immunofluorescence of H3K27ac (green, using anti-H3K27ac antibody) and NHA9 labelled by LD655 via OTO-GCE (red) to assay their colocalisation. Right: Line scans along the white lines in the magnified images in left. **d** Schematic of the FLIM-FRET analysis pipeline. NHA9 condensates were labelled by FRET dye pair via OTO-GCE, followed by acquiring donor decay profiles cell-by-cell. Nuclear condensates were selected as a region of interest to extract donor decay profiles afterwards. **e** Acceptor photobleaching assays for both single-mutant and double-mutant NHA9 labelled with FRET dye pair. The averaged donor lifetime distributions were acquired before and after Acceptor photobleaching to infer FRET events exist or not. **f** Normalised donor fluorescence decays of five NHA9 variants labelled by FRET dye pair in cells. **g** *R_E_* extracted from lifetime-based fitting and scaling law of the NHA9 FG domain in cells from global fitting. Scale bars: 5 µm.

To confirm selective labelling of NHA9 by the OTO-GCE system, cells were co-transfected with plasmids encoding amber-mutated NHA9 and wild-type NHA9 fused to mEGFP. Upon treatment with TCO*A and LD655-H-tetrazine, colocalisation of LD655 and mEGFP-NHA9 signals was observed via fluorescence microscopy (Fig. 5b). Additionally, immunofluorescence analysis confirmed the colocalisation of NHA9 condensates with histone H3 lysine 27 acetylation (H3K27ac), consistent with previous reports^29^ (Fig. 5c). These results verify the presence of TCO*A and that the fluorophore was selectively incorporated into NHA9 condensates.

To accurately assess the conformational dimensions of the NHA9 FG domain, intermolecular FRET between different NHA9 molecules needed to be excluded. This was achieved by co-transfecting cells with a mixture of wild-type and TAG-mutated NHA9 plasmids to ensure a low concentration of labelled protein within condensates (Supplementary Fig. 8). To further confirm the absence of intermolecular FRET, we expressed NHA9_A221TAG_ in combination with wild-type NHA9, followed by treatment with a mixture of donor and acceptor dyes. As a result, each NHA9 molecule could be labelled with either a donor or an acceptor dye, but not both. Condensates were selected based on low acceptor intensity per pixel (excited at 660 nm) to minimise the likelihood of intermolecular FRET (Supplementary Fig. 9b). Donor lifetimes were measured before and after acceptor photobleaching. An increase in donor lifetime post-photobleaching would indicate intermolecular FRET, whereas unchanged lifetimes would confirm its absence. We observed no change in donor fluorescence lifetime after acceptor photobleaching, indicating that intermolecular FRET did not occur (Fig. 5d,e).

We examined five dual-labelled FG-domain variants within cellular NHA9 condensates. Data were collected from around 30 cells per variant to account for cellular heterogeneity. Analysis of donor fluorescence lifetime decay curves revealed differences in FRET efficiency among the variants (Fig. 5f). These differences were further visualised using phasor plot analysis (Supplementary Fig. 9c), which confirmed the observed trend. To extract the apparent Flory scaling exponent (ν), we performed global fitting of the decay curves using the Gaussian chain model according to Eq. 10 (see “Methods” for details), yielding ν = 0.61 ± 0.01 (Fig. 5g). Additionally, a mutant-by-mutant analysis without assuming a specific model yielded ν = 0.63 ± 0.25, in good agreement with the global fit (Supplementary Fig. 9d).

### Molecular simulations confirm NHA9’s IDP expansion within nanocluster and reveal micelle-like structures

To obtain a more detailed understanding of protein conformations, we performed molecular dynamics simulations of NHA9. Residue-level coarse-grained simulations have proven effective in describing protein phase behavior and capturing trends in experimental phase separation propensities. We employed the HPS-Urry force field, a model specifically developed for implicit solvent simulations of IDPs^55^. The secondary and tertiary structures of HOXA9 were constrained using an elastic network model. Simulations were conducted with varying numbers of NHA9 chains (ranging from 1 to 50) to investigate individual protein conformations across clusters of different sizes. We calculated the mean distances (*R*_E_) between residues which were fluorescently labelled in the FRET experiments (Fig. 6a). These labelled residues, all located within the disordered FG domain of NHA9, exhibited trends consistent with experimental observations: Single NHA9 chains adopted highly collapsed conformations, whereas inter-residue distances increased with cluster size, indicating protein expansion the nanocluster and condensate regimes. This behavior is consistent with that observed for other IDPs^56^.

**Figure 6.**
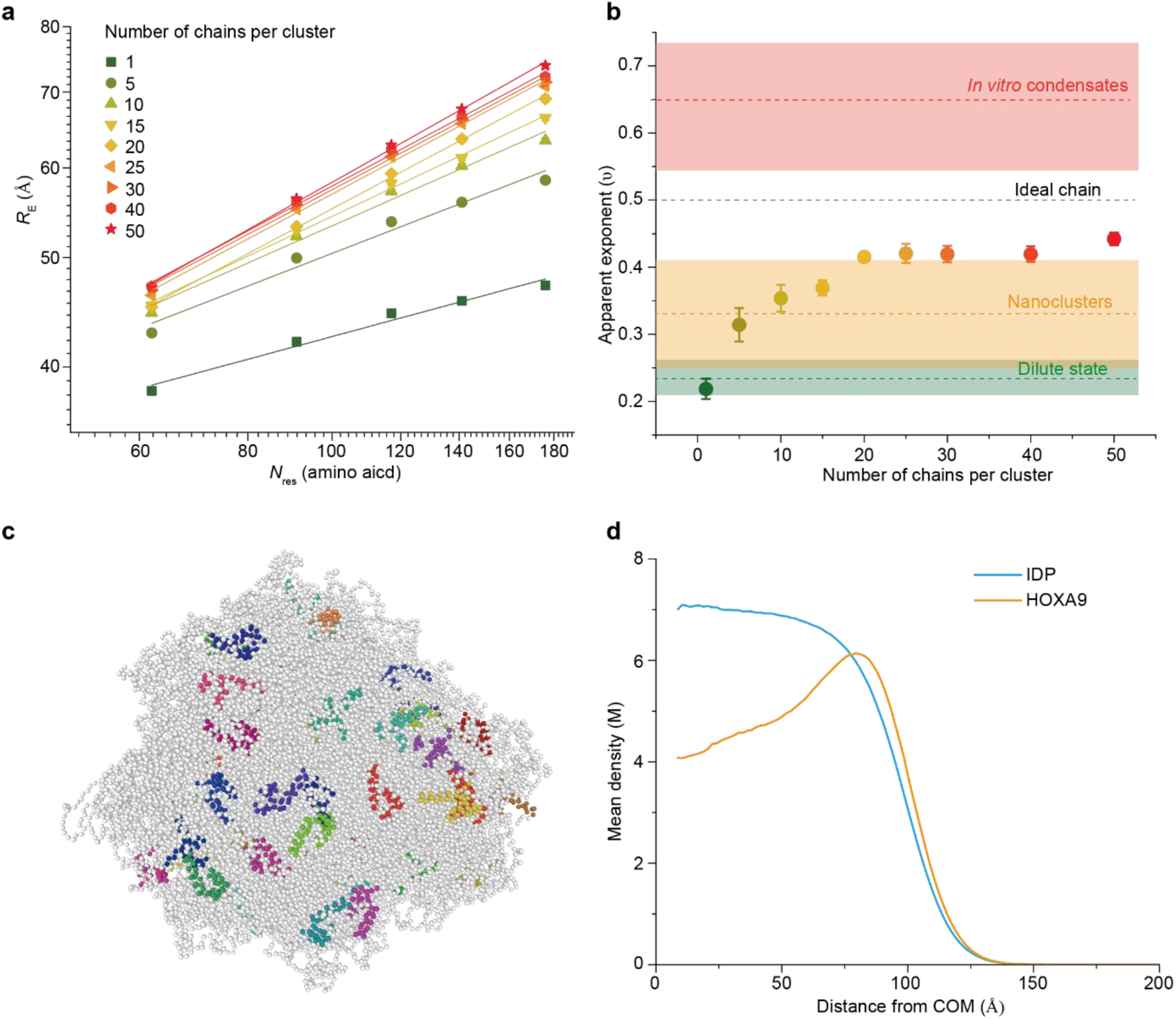
Coarse-grained simulations of NHA9 in dilute states and nanocluster. **a** Mean end-to-end distances (*R*_E_) vs sequence distance (*N*_res_) for clusters of different sizes extracted from coarse-grained simulations. The solid lines show scaling law fitting. **b** Apparent Flory exponent (ν) as a function of the number of chains in cluster, obtained by fitting the data points in shown in Fig. 6a. Data are presented as mean +/− residual standard deviation. Dashed lines represent scaling exponent values derived from experimental measurements across different assembly states. Colored rectangles represent fitted scaling values +/− standard error of the estimate from experimental measurements. **c** Snapshot of a cluster containing 50 NHA9 chains. The colored amino acids represent the helices in the DBD and the remaining amino acids are shown in white. Snapshot was made with OVITO^80^. **d** Mean density of each block in NHA9 as a function of its distance from the cluster’s center of mass. Results for the cluster contains 40 chains.

By fitting the data presented in Figure 6a, we computed the apparent Flory exponents for our simulated systems (Fig. 6b). For the single-chain system, we obtained a scaling exponent of ν = 0.21 ± 0.02, closely matching the experimentally measured value of ν = 0.24 ± 0.02. For cluster sizes of 5 and 10 chains, the exponents increased to ν = 0.31 ± 0.02 and ν = 0.35 ± 0.02, respectively. Beyond these sizes, ν continued to rise, reaching a value of ν = 0.44 ± 0.01 for the largest simulated cluster of 50 chains. These values are consistent with those obtained experimentally in the subsaturated regime, where nanoclusters form (ν = 0.33 ± 0.08). Due to computational limitations, we were unable to simulate sufficiently large systems to observe further expanded IDPs. Moreover, stretching of the shortest segment involved in the FRET measurements (62 aa) appeared to saturate at a cluster size of 30 chains, whereas longer segments continued to exhibit some degree of expansion as the cluster size increased from 30 to 50 chains. This suggests that further increases in cluster size would lead to further expansion of the IDP block, and increase the observed value of ν. The heteropolymer nature of proteins may play a role here, as evidenced by heterogeneous structures recently reported in condensates as well as the approximate nature of coarse-grained force-fields^57^.

Our simulations captured the expansion behaviour of the IDP block of NHA9 in line with the available experimental data, therefore we aimed to extend our analysis to the behaviour of the HOXA9 block. Snapshots from the simulated trajectories clearly show that the structured DBD preferentially localises near the surface of the nanoclusters (Fig. 6c, Supplementary Fig. 10). This localisation can be explain by the high net positive charge of the DBD (+8e, or +0.14e per residue), rendering it highly hydrophilic and leading to repulsive interactions with the hydrophobic IDP block, which also carries a slight positive charge (+0.02e per residue). To quantify this localisation bias, we computed density profiles as a function of distance from the cluster’s center of mass (COM) for both the IDP and HOXA9 blocks of NHA9 (Fig. 6d, Supplementary Fig. 11 and 12). The results show that IDP residues are enriched near the cluster’s center, whereas the HOXA9 block exhibits a strong preference for localization near the interface, resembling micelle-like structures. This behavior aligns with the observed differences in hydrophobicity and net charge between the two domains and suggests a propensity for microstructure formation within the clusters. Additionally, the FG2 domain was preferentially located near the center of the condensate, while the FG1 domain, positioned at the chain end, was enriched at the interface (Supplementary Fig. 13 and Fig. 14), consistent with previous observations in polymer droplets^58^. This segregation becomes less pronounced as the cluster grows and both FG domains migrate to the center of the condensate. Our findings also indicate that the DBD has a strong propensity to be at least partially in contact with the solvent, suggesting the preservation of its native structure, which allows for DNA binding at the surface of the clusters. We also computed the intermolecular residue-residue contact maps, as shown in Supplementary Fig. 15. The analysis revealed an extensive network of contacts between the mostly hydrophobic residues in the IDP. This suggests that nanocluster formation is mediated mostly by hydrophobic IDP-IDP interactions, whereas the HOXA9 block forms, on average, fewer and more specific contacts.

## Discussion

Transcriptional condensates play a critical role in regulating gene expression by organising transcription factors and the associated transcription machinery into functional compartments^7,11,59^. Increasing evidence indicates that many proteins prone to condensation are associated with the early formation of large nanoclusters, both below and above their phase separation thresholds^32,34^. In this study, we show that NHA9 forms nanoclusters at subsaturated concentrations, below the critical concentration for phase separation. Furthermore, we demonstrate that DNA binding modulates NHA9 nanocluster formation, suggesting a regulatory mechanism in which chromatin interactions can influence protein assembly dynamics. Given that NHA9 nanoclusters arise at concentrations below 100 nM, it is likely that they form under physiological conditions and contribute to transcriptional regulation through a mechanism distinct from classical phase-separated condensates, similar to the reported gene-regulatory role of the transcription factor Mig1 clusters^35^.

The distinct assembly states of NHA9 are primarily driven by its long, intrinsically disordered FG domain. However, the precise molecular mechanisms underlying NHA9 self-assembly and its structural dynamics remain poorly understood. To elucidate these mechanisms, we systematically investigated NHA9 behaviour across a range of protein concentrations. Our data revealed a conformational transition in the FG domain, progressing from a dilute monomeric state to an oligomeric state, and ultimately to a dense phase (Fig. 7a, b). Both inter- and intramolecular interactions mediate these transitions. In the dense phase, extensive protein-protein interactions promote chain expansion, whereas in the dilute phase, intramolecular contacts dominate, resulting in a more compact conformation that minimises free energy. Surprisingly, our results suggest that the oligomeric state is structurally similar to micelles, with the hydrophilic DBD domain located near the surface—a feature also predicted by computational simulations for Nanog, a master TF^60^. Experimentally, the number of chains in the structure increases with overall concentration, hinting at morphologies that are not typical core-shell micelles. The NHA9 micelles have different appearances from those micelles recently identified for TDP43 and SRSF proteins, which present fixed-size morphologies on electron micrographs^61^. NHA9 exhibits a weaker amphiphilic block copolymer character compared to TDP-43 and SRSF, potentially explaining the observed differences in micellar appearance. Our simulation results nevertheless indicate that NHA9 nanoclusters possess a distinct substructure wherein the DBD is oriented toward the exterior despite their inherent heterogeneity and stochastic assembly. This structural feature appears intrinsically encoded within the oncofusion block copolymer NHA9 sequence properties. Unlike typical surfactants, we observe phase separation at sufficiently high concentrations. In block copolymers, a competition between phase separation and micellisation under weakly amphiphilic conditions can drive transitions between micellar and phase-separated regimes^62,63^. Consequently, the presence of phase separation at higher concentrations does not preclude the formation of biologically functional micelles at low concentrations.

**Figure 7.**
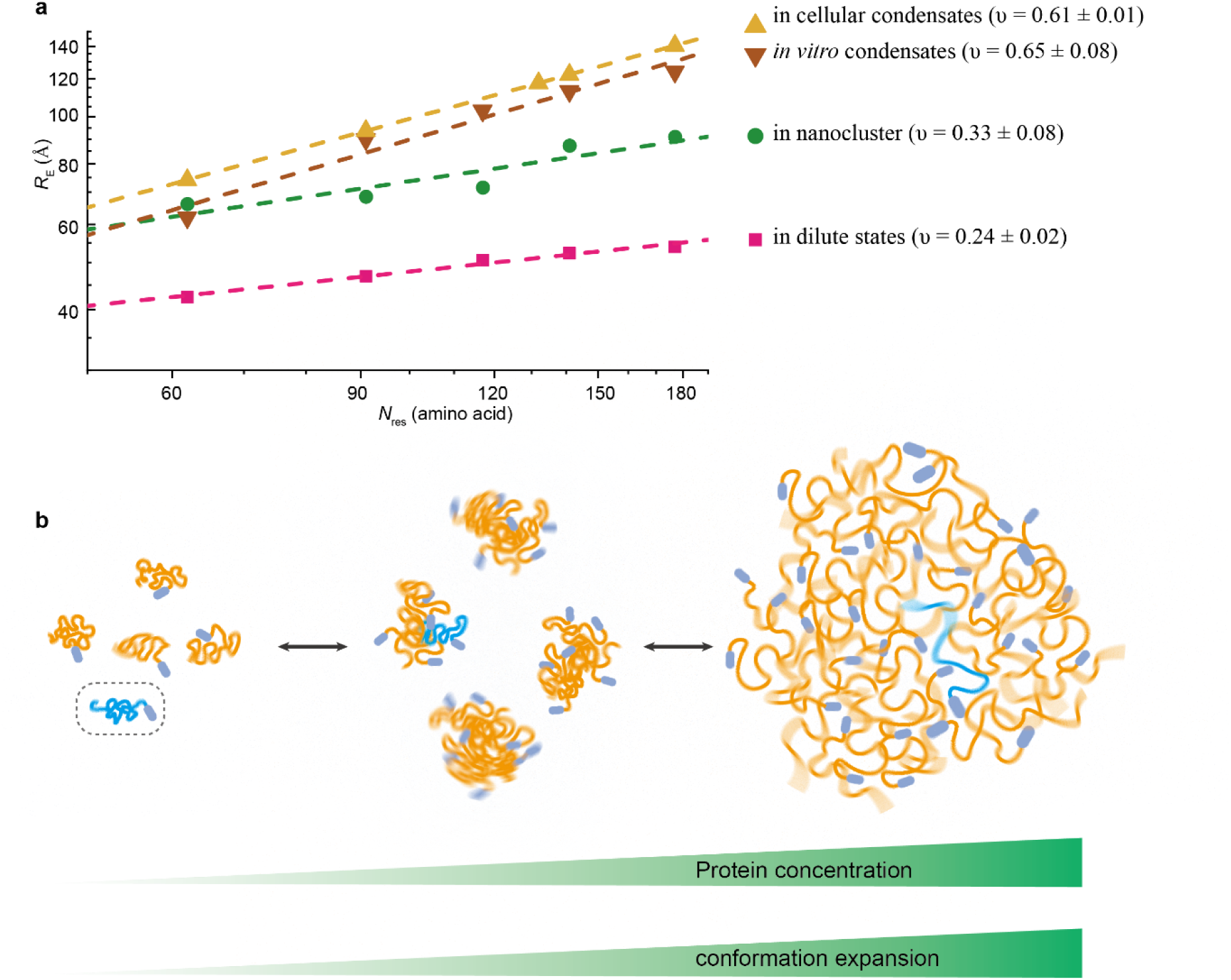
Concentration dependent self-assembly of NHA9. **a** Summary of scaling law of FG domain of NHA9 under different assembly states. The disordered FG domain is the driving force of NHA9 self-assembly and our data revealed that the FG domain became more and more expanded from the dilute state, to nanocluster and condensates. **b** Conceptual scheme illustrating NHA9 FG domain expansion within different assembly state.

In the cellular environment, transcription factor condensates function as dynamic hubs that recruit components of the transcriptional machinery—including mediator, general transcription factors, and RNA polymerase II—to facilitate transcription initiation. Our *in situ* measurements reveal that the FG domain of NHA9 adopts a more extended conformation in cellular phase-separated condensates (ν = 0.61 ± 0.01), which is consistent with *in vitro* condensates (ν = 0.65 ± 0.08). This conformation results from self-association and may also enhance interactions between NHA9 and components of the transcription machinery. While the cellular FLIM-FRET measurements provided valuable insights, they are not without limitations that may influence the results. First, the data were obtained from transiently transfected cells, which results in NHA9 overexpression and does not reflect endogenous expression levels. Consequently, the observed NHA9 condensates often exceed 1 μm in diameter, potentially diverging from physiological conditions. TFs expressed at endogenous levels generally form numerous smaller transcriptional hubs or condensates; for example, HSF1 forms condensates with diameters of approximately 300 nm^10,18^. Recent studies have further shown that such localised, high-concentration transcriptional hubs—typically containing fewer than 100 molecules and exhibiting dwell times of approximately 1–2 minutes—are critical for effective transcriptional activation^64^. Furthermore, smaller condensates with signals near background levels were excluded in our FLIM-FRET analyses, limiting our observations to larger structures. Second, the GCE approach used for site-specific labelling introduces an in-frame stop codon, potentially resulting in truncated NHA9 variants that may influence condensate behavior. Third, although mild plasma membrane permeabilisation facilitated the use of hydrophilic dyes with reduced nonspecific binding, some background signal from free dyes likely remained, potentially reducing the signal-to-noise ratio. Future studies should focus on optimising GCE strategies to prevent protein truncation and enable endogenous-level labelling and on employing fluorogenic, cell-permeable dyes to enhance imaging specificity and quality, which might allow the study of conformational ensembles of nanoclusters in the future. Despite these technical constraints, our *in situ* scaling measurements avoided using large fluorescent label, such as GFP or self-labelling proteins, thereby minimizing potential perturbation of NHA9 condensates in cells. Most importantly, the data support the conclusion that the disordered FG domain of NHA9 adopts a similarly extended conformation in both *in vitro* and cellular environments.

Our study provides a molecular-level understanding of biomolecular conformational shifts during the condensation process. At low concentrations, proteins exist as individual molecules in the dilute phase. As concentration increases, they begin assembling into nanoclusters, and above the saturation threshold, these clusters coalesce into a condensed phase. Importantly, our findings reveal that this assembly process is accompanied by a continuous conformational transition from a compact chain in the monomeric state to expanded ones in the oligomeric state, culminating in more stretched chains in a densely packed phase-separated condensate. Furthermore, we found that the oncofusion protein NHA9 exhibits characteristics analogous to a weakly amphiphilic diblock polymer. Specifically, the disordered FG-rich domain in NHA9 resembles a hydrophobic block, while the DBD resembles a hydrophilic block. Notably, this weak amphiphilic architecture enables NHA9 to self-assemble into non–core-shell micelles, with the DBD preferentially oriented towards the exterior. Such a configuration is likely functionally significant, as it facilitates direct access of DNA to the DBD, thereby promoting DNA interactions, which could also orchestrate long-range chromatin interactions such as enhancer-promoter looping at oncogene loci. Since oncofusion TFs are typically formed from two blocks, an IDP block and a DBD block, their block copolymer–like properties may represent a common feature among oncofusion TFs. The formation of non–core-shell spherical micelles could serve as a general assembly principle underlying their oncogenic activity. As many eukaryotic TFs already obey blockcopolymer charateristics, we could expect formation of non–core-shell spherical micelles also for those with weak amphiphilicity and a long IDP region. These insights contribute to a deeper understanding of the molecular principles governing transcriptional condensates and highlight the importance of structural transitions in regulating transcriptional activity.

## Methods

### Protein expression and purification

The human NHA9 protein lacking the Gle2-binding sequence (GLEBS; amino acids 157–203) was cloned into the bacterial expression vector pQE-14His-TEV. Recombinant NHA9 was expressed in *Escherichia coli* strain BL21-AI. Cultures were grown in Luria-Bertani (LB) medium supplemented with 50 μg/mL kanamycin at 37 °C until reaching an optical density at 600 nm (OD₆₀₀) of approximately 0.6. Protein expression was induced with 1 mM isopropyl-β-d-thiogalactoside (IPTG) and 0.02% (w/v) arabinose at 18 °C. Following overnight expression, cells were harvested by centrifugation and resuspended in lysis buffer containing 6 M guanidine hydrochloride, 50 mM Tris-HCl, 1 M NaCl, 20 mM imidazole, and 2 mM 2-mercaptoethanol at pH 9.0. Cells were lysed by sonication, and the lysate was clarified by centrifugation at 19,000 × g for 1 h at 4 °C. The supernatant was incubated with Ni-NTA agarose beads for 2 h at 4 °C. Beads were subsequently washed with buffer containing 4 M guanidine hydrochloride, 50 mM Tris-HCl, 2 M NaCl, 40 mM imidazole, and 2 mM 2-mercaptoethanol at pH 9.0. The his-tagged protein was eluted with buffer containing 2 M guanidine hydrochloride and 50 mM Tris-HCl at pH 8.0. Eluted proteins were dialysed against buffer containing 0.5 M guanidine hydrochloride and 50 mM Tris-HCl at pH 8.0 using dialysis membranes with a molecular weight cutoff (MWCO) of 6–8 kDa to remove imidazole. His-tags were cleaved overnight at room temperature with TEV protease. Typically, proteins precipitated following cleavage and were resolubilised in 4 M guanidine hydrochloride at pH 9.0. The solution was incubated again with Ni-NTA beads to remove uncleaved protein and the cleaved his-tag. The flow-through was concentrated and further purified by size-exclusion chromatography using a Superdex 75 column equilibrated with 4 M guanidine hydrochloride, 50 mM Tris-HCl, 2 M NaCl, and 0.2 mM TCEP at pH 9.0. Fractions were analysed by SDS-PAGE using 4–12% gradient gels with MOPS running buffer and stained with Coomassie Brilliant Blue. Pure fractions were pooled and concentrated to approximately 15 mg/mL using 3-kDa MWCO centrifugal filters (Merck Millipore), and protein concentrations were determined using the BCA Protein Assay Kit (Thermo Fisher). Final protein preparations were flash-frozen in liquid nitrogen and stored at −80 °C.

For constructs containing a TGT codon (encoding cysteine) at the second residue and a TAG amber codon at position A221 (to incorporate p-acetylphenylalanine, AcF, via amber suppression), sequences were cloned into the pQE-14His-TEV vector. All constructs were co-transformed with the pEvol plasmid, encoding an evolved aminoacyl-tRNA synthetase specific for AcF and its cognate tRNA, into *Escherichia coli* BL21-AI cells^65^. Cultures were grown in LB medium supplemented with 50 μg/mL kanamycin and 33 μg/mL chloramphenicol at 37 °C until OD₆₀₀ reached 0.2, at which point 1 mM AcF was added. Induction and purification followed the same procedure as described for the non-TAG constructs.

### NHA9 FG domain labelling *in vitro*

#### NHA9 labelling for FLIM-FRET measurements in reconstituted condensates

Purified NHA9 containing double cysteine mutations was reduced with 10 mM 1,4-dithiothreitol (DTT) at room temperature (RT) prior to labelling with thiol-reactive dyes. Following reduction, the protein was buffer-exchanged into maleimide labelling buffer (4 M guanidine hydrochloride, 1× PBS, 0.1 mM EDTA, 0.2 mM TCEP, pH 7.0) using a centrifugal filter with a 3 kDa molecular weight cutoff (MWCO; Amicon). Labelling was performed overnight at 4 °C using Alexa Fluor 594 maleimide (A10256, Thermo Fisher) and LD655-maleimide (Lumidyne Technologies) at a 1:2 dye-to-protein molar ratio. The reaction was quenched by adding DTT to a final concentration of 10 mM. Free dyes were removed via buffer exchange followed by gel filtration using a Superdex 75 column.

#### NHA9 labelling for smFRET measurements

For single-molecule FRET (smFRET) labelling, NHA9 variants containing a single cysteine and a TAG amber codon were site-specifically labelled with Alexa Fluor 594-maleimide at the cysteine residue (Acceptor) and Alexa Fluor 488-hydroxylamine at the AcF residue (donor), following established protocols^65^. Briefly, samples were exchanged into oxime labelling buffer (50 mM sodium acetate-HCl, pH 4.0, 150 mM NaCl, 4 M guanidine hydrochloride, 100 mM aniline). A total of 50 µL of 200 µM protein was reacted with 50 µL of 1 mM Alexa Fluor 488-hydroxylamine (a fivefold molar excess of dye) for 24 h at 37 °C. After the reaction, samples were exchanged into maleimide labelling buffer through three rounds of buffer exchange using a 3 kDa MWCO centrifugal filter (Amicon), followed by reduction with 10 mM DTT. DTT was then removed by five additional rounds of buffer exchange. The freshly prepared protein was subsequently labelled with Alexa Fluor 594 maleimide (1:2 dye-to-protein molar ratio) overnight at 4 °C. The reaction was quenched with 10 mM DTT, and unreacted dye was removed by buffer exchange and gel filtration using a Superdex 75 column.

### Electrophoretic mobility shift assay (EMSA)

Varying concentrations of NHA9 (up to 1 µM) were incubated with 10 nM Cy5-labelled double-stranded DNA (5’-ACTCTATGATTTACGACGCT-3’ as previously reported^30^; HOXA9 binding site, TTTAC) in a buffer containing 50 mM Tris-HCl (pH 7.4), 150 mM NaCl, 1 mM EDTA, 0.01% (v/v) Tween-20, and 3% glycerol. After mixing, the protein-DNA complexes were incubated for 5 min at room temperature. Samples were then resolved on a 6% native polyacrylamide gel prepared in 1× TBE buffer and electrophoresed at a constant voltage of 100 V for 40 min. Fluorescently labelled DNA-protein complexes were visualised using a ChemiDoc MP imaging system (Bio-Rad).

### Turbidity measurement

Purified NHA9 in 2 M GdmCl was rapidly mixed with 24 μL of buffer (50 mM Tris-HCl, pH 7.4, 150 mM NaCl). The resulting mixture (total volume: 25 μL) was immediately transferred to a clear-bottom 384-well plate, and turbidity was assessed by measuring absorbance at 340 nm using a SpectraMax iD5 plate reader. All measurements were performed in triplicate for each construct under the specified conditions.

### Cell culture, transient transfection and labelling

The HeLa Kyoto cell line (RRID: CVCL_1922), kindly provided by Martin Beck’s laboratory, was cultured in DMEM supplemented with 1 g/L glucose (Gibco, 31885023), 9% fetal bovine serum (FBS; Sigma, F7524), 1% penicillin-streptomycin (Thermo Fisher, 15140-122), and 1% L-glutamine (Thermo Fisher, 25030-081) at 37 °C in a humidified incubator with 5% CO₂. Cells were routinely tested and confirmed to be free of mycoplasma contamination.

To introduce NHA9 transcriptional condensates, HeLa cells were transiently transfected with plasmids encoding NHA9. Co-transfection with plasmids encoding TAG-mutated NHA9 and the OTO-GCE system enabled site-specific labelling. To minimise intermolecular FRET signals caused by heterogeneity of transient expression, wild-type NHA9 was co-expressed in excess. For this purpose, a bicistronic construct (pBI_Flag::NHA9TAG::HA(boxB)_Flag::NHA9::HA::P2A-T2A::NHA9) was designed using the pBI_CMV vector and self-cleaving peptides to promote higher expression of wild-type NHA9 relative to the TAG-mutant. Additional plasmids encoding wild-type NHA9 were also co-transfected to further increase the expression ratio. For FLIM-FRET experiments, 150,000 HeLa cells were seeded per 35 mm µ-dish (ibidi, 81158) one day prior to transfection. Transfection was performed using jetPRIME reagent (Polyplus-transfection) with 1.8 µg total DNA, 150 µL jetPRIME buffer, and 4 µL jetPRIME reagent per dish. A 1:1:1 mass ratio of the following plasmids was used:

- pBI_Flag::NHA9::HA
- pBI_Flag::NHA9^TAG^::HA(boxB)_Flag::NHA9::HA::P2AT2A::NHA9
- pcDNA3.1_TOM20_1-70_::FUS_1-478_::V5::Myc::4xλ_N22_::NES::PylRS^Y306A,Y384F^_U6-tRNA^Pyl,CUA^

A list of all plasmids used is provided in Supplementary Table 2. Four to five hours post-transfection, the culture medium was replaced with fresh medium containing 60 μM trans-cyclooct-2-en-l-lysine (TCO*A; SciChem) for in-cell labelling, or tert-butyloxycarbonyl-l-lysine (BOC; Iris Biotech) for control experiments. Media were also supplemented with 10 mM HEPES (pH 7.25). Stock solutions of 100 mM for all ncAAs were prepared in 15% (v/v) DMSO/0.2 M NaOH, following established protocols^52^.

Twenty to twenty-four hours after transfection, cells were washed twice with transport buffer (TB: 20 mM HEPES, 110 mM KOAc, 5 mM NaOAc, 2 mM MgOAc, 1 mM EGTA, 2 mM DTT, pH 7.3 adjusted with KOH), supplemented with 5 mg/mL PEG6000 to prevent osmotic shock. To minimise interference from cellular autofluorescence in the green spectral range, we employed the AF594-H-tetrazine and LD655-H-tetrazine dye pair. Cellular delivery of the dyes was achieved via mild digitonin treatment, which permeabilises the plasma membrane while preserving nuclear envelope integrity. Specifically, cells were then permeabilised by incubation with 40 µg/mL digitonin (AppliChem, A1905) for 10 min at room temperature. Following two washes with TB, cells were incubated in a dye solution containing 33.3 nM Alexa Fluor 594-H-tetrazine (Click Chemistry Tools) and 66.6 nM LD655-H-tetrazine (Lumidyne Technologies) in TB for 5 min at room temperature. A 1:2 donor-to-acceptor dye ratio was used to reduce the donor-only population^49^. After labelling, cells were washed in TB and incubated at 37 °C for 30 min to remove residual dyes. Cells were used for imaging within a 3-hour window post-labelling.

### Western blot assay

HeLa cells were seeded in a 6-well plate 15–20 h prior to transfection to reach 70–80% confluency at the time of transfection. Each well was transfected with 3 μg of the plasmid [pBI_Flag::NHA9TAG::HA(boxB)_Flag::NHA9::HA::P2A-T2A::NHA9] using jetPRIME (Polyplus-transfection). Twenty to twenty-four hours after medium replacement, cells were harvested and lysed in 40 μL RIPA buffer (150 mM NaCl, 1.0% Triton X-100, 0.5% sodium deoxycholate, 0.1% SDS, 50 mM Tris-HCl, pH 8.0) supplemented with Complete Protease Inhibitor Cocktail (Roche, 11873580001), 1 mM MgCl₂, and Sm nuclease (Protein Production Core Facility, IMB Mainz). Protein lysates were resolved by SDS-PAGE using NuPAGE 4–12% Bis-Tris gels with NuPAGE MOPS SDS Running Buffer (Thermo Fisher Scientific), and transferred to nitrocellulose membranes (0.2 µm pore size; Trans-Blot Turbo Midi, Bio-Rad, 1704159) using the Trans-Blot Turbo Transfer System (Bio-Rad). Membranes were blocked for 1 hour at room temperature (RT) in 5% low-fat milk (Carl Roth, T145.3) prepared in PBS. Blots were incubated overnight at 4 °C with a mouse anti-HA primary antibody (2 μg/mL; Protein Production Core Facility, IMB Mainz) in blocking solution. The following day, membranes were washed with PBST (PBS containing 0.2% Tween-20), then incubated with a secondary antibody (goat anti-mouse HRP-conjugated, 1:10,000 dilution; Jackson ImmunoResearch, 715-035-150) for 1 hour at RT. After final washes with PBST, membranes were incubated for 1 min with enhanced chemiluminescence substrate (ECL; Cytiva, RPN2106). Signal detection was performed using a ChemiDoc MP imaging system (Bio-Rad).

### Fixed cell immunofluorescence

HeLa cells were transfected with the pcDNA3.1_mEGFP::4×GGS::NHA9 plasmid (where 4×GGS serves as a flexible linker) and incubated for 24 h prior to fixation. Cells were fixed with 2% paraformaldehyde in PBS for 10 min at room temperature (RT), permeabilised with 0.5% Triton X-100 in PBS for 15 min at RT, and rinsed twice with PBS before blocking. Blocking was performed using 3% bovine serum albumin (BSA) in PBS for 90 min at RT. Cells were then incubated overnight at 4 °C with a primary antibody against H3K27Ac (histone H3 lysine 27 acetylation; 1:1000 dilution, Abcam, ab4729) diluted in blocking buffer. The following day, samples were washed with PBS and incubated with a secondary antibody (1:2000 dilution, Abcam, ab205722) for 60 min at RT. After final rinses with PBS, fresh PBS was added to the wells, and cells were prepared for imaging.

### Single-molecule FRET measurements

Single-molecule fluorescence experiments were conducted using a custom-built multiparameter spectrometer centered around a high-numerical-aperture water-immersion objective (Nikon 60×, 1.27 NA) mounted on a z-translator, as previously described^40^. Briefly, linearly polarised outputs from three picosecond pulsed laser diodes—485 nm (LDH-D-C-485, PicoQuant), 560 nm (LDH-D-TA-560, PicoQuant), and 660 nm (LDH-D-C-660, PicoQuant)—were passed through corresponding excitation filters (488/10, 560/14, and 661/11). Fluorescence emission was spatially filtered using a 100 µm pinhole, then split into parallel and perpendicular polarisation components. Each polarisation component was further spectrally separated into green, orange, and red channels using emission filters (525/50, 609/57, and 700/75, respectively) and focused onto single-photon counting detectors (green: MPD, PicoQuant; orange: PMA Hybrid 40, PicoQuant; red: τ-SPAD, PicoQuant). Laser pulses were alternated to probe the presence of acceptor fluorophores, with synchronisation controlled by a multichannel laser driver (Sepia II, PDL 828, PicoQuant)^66,67^. The entire setup was operated using SymPhoTime64 software (PicoQuant GmbH). Samples labelled with Alexa Fluor 488 and Alexa Fluor 594 were excited using the 485 nm laser at 55 µW and the 560 nm laser at 65 µW, both operating at a repetition rate of each 25 MHz. Measurements were performed at a sample concentration of ∼50 pM in buffer containing 50 mM Tris-HCl (pH 7.4), 150 mM NaCl, 0.01% Tween-20, and freshly added 10 mM DTT. Photon detection was performed using a multichannel time-correlated single-photon counting module (HydraHarp 400, PicoQuant) with a time resolution of 16 ps. Polarisation correction factors were determined to be *L*_1_ = 0.16 and *L*_2_ = 0.

### Analysis of transfer efficiency histograms

Single-molecule fluorescence data were analysed using the PAM software ^68^. Single-molecule events were identified using the All-Photon Burst Search algorithm with a threshold of 5 photons per 1000 µs time window and a minimum total photon count of 50 per burst. The FRET transfer efficiency (*E*) and stoichiometry (*S)* for each burst were calculated as follows:

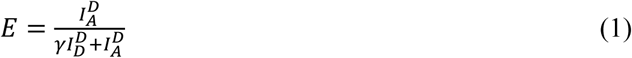

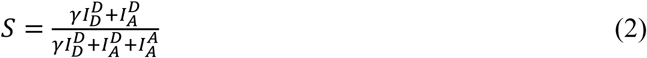

where 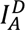 is the corrected intensity from the Acceptor after donor excitation, 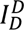 is the corrected intensity from the donor after donor excitation and 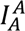 is the corrected intensity from the Acceptor after acceptor excitation. Raw intensities were corrected for background, donor signal leakage into the acceptor channel (α = 0.086), and direct excitation of the Acceptor by the donor-exciting laser (δ = 0.046). The γ factor accounts for differences in quantum yield and detection efficiency between the donor and acceptor fluorophores (γ = 0.323).

To estimate the mean transfer efficiency, the transfer efficiency histograms were fitted with a 2D Gaussian function. For smFRET measurements of nanoclusters, stoichiometry values of nanocluster species were below 0.5 because of acceptor quenching. The Donor-only species and FRET species, indicated by blue rectangles, were individually selected for separate fitting using a 2D Gaussian function, as shown in Supplementary Figure 3e. The fitted mean transfer efficiencies ⟨*E*⟩ values were applied to acquire the root-mean-square end-to-end distances 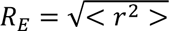 by numerically solving the following equation:

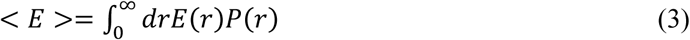

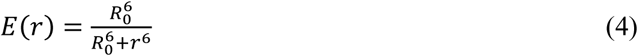

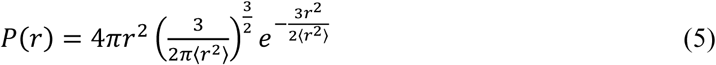

where *R*_0_ is the Fӧrster radius with *R*_0_ = 5.6 nm for a Alexa488/Alexa594 dye pair and *R*_0_ = 7.7 nm for a Alexa594/LD655 dye pair according to previous reports^40,42^. The distance distribution function between the donor and acceptor dyes, *P*(*r*), we used a Gaussian chain model, given by Eq. 5. In smFRET measurements the donor and acceptor anisotropies are low (*r*<0.3), indicating the free rotation of dyes. Finally, the extracted *R*_E_ values were used to determine the polymer scaling law factor by fitting to:

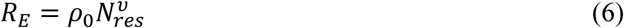

### Fluorescence correlation spectroscopy (FCS) measurements

FCS measurements were performed using the same setup as for smFRET experiments, employing a 660 nm laser (LDH-D-C-660, PicoQuant) operated in continuous-wave mode at a power of 50 μW with a 50 μm pinhole. Experiments were conducted with 10 nM LD655-labelled NHA9 protein mixed with varying concentrations of unlabelled NHA9 in buffer containing 50 mM Tris-HCl (pH 7.4), 150 mM NaCl, 0.01% Tween-20, and freshly added 10 mM DTT. A total of 120 measurements, each with a 10-second acquisition time, were collected for each condition. Data analysis was carried out using SymPhoTime 64 software. Autocorrelation curves were generated for lag times ranging from 0.0001 ms to 1000 ms and fitted using a standard diffusion model (Eq. 7), where *ω*_0_⁄*ω*_*Z*_ is structure parameter, *τ*_*D*_ and *N* represent the readout parameters of the fit.

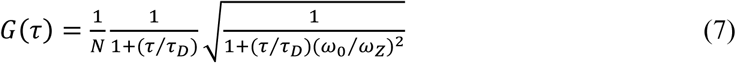

### *In vitro* droplet assay and FLIM–FRET measurements

Purified labelled and unlabelled NHA9 proteins were mixed at a 1:10,000 ratio in 2 M guanidine hydrochloride. A 1 μL aliquot of the mixed protein solution was rapidly diluted into 24 μL of assay buffer (50 mM Tris-HCl, 150 mM NaCl, 10 mM DTT) on a 15-well µ-slide (ibidi, 81507, Germany). The final concentrations were 10 μM unlabelled and 1 nM labelled NHA9. FLIM acquisition was initiated within the first 5 min after forming the droplets, as the condensates were more liquid during this time. Over time the droplets hardened out, likely due to molecular ageing as observed also for Nup98 FG domains^69^. Samples were excited using a pulsed laser operating at 40 MHz with a power of 70 μW. Imaging was performed using SymPhoTime 64 software with the following parameters: pixel size of 200 nm, dwell time of 100 μs, image resolution of 256 × 256 pixels, and a time resolution of 16 ps. Fluorescence anisotropy values are presented in Supplementary Figure 6a and 6b.

### Fluorescence recovery after photo bleaching (FRAP)

NHA9 droplets were prepared as described above. FRAP experiments were performed on a Visitron spinning-disk microscope (Visitron Systems GmbH) together with an ORCA-flash 4.0 camera C13440 CMOS camera (Hamamatsu Photonics). Bleaching was done with a 488 nm laser at 70% power. Timelapse images were acquired over a 98 s time course after bleaching with a 2 s interval. Fluorescence intensities of regions of interest were corrected by unbleached control regions and then normalized to pre-bleached intensities of the regions of interest.

### FLIM-FRET for cell measurements

The same optical setup used for smFRET was employed for FLIM-FRET measurements, both *in vitro* (droplets) and in live cells. For cellular imaging, the following acquisition parameters were used: pixel size of 100 nm, image resolution of 256 × 256 pixels, pixel dwell time of 150 μs, and time resolution of 16 ps. To ensure proper sample quality, labelled cells were first imaged under 660 nm laser excitation to confirm the absence of cytoplasmic aggregates (“blobs”), which indicate the presence of insoluble NHA9 truncation products. The acceptor fluorescence intensity per pixel within nuclear condensates was also assessed under 660 nm excitation to evaluate expression levels and avoid overexpression, which could result in intermolecular FRET. Expression levels were further validated through acceptor photobleaching. A constant donor fluorescence lifetime before and after photobleaching confirmed the absence of intermolecular FRET (see Figure 5e). Cells that met these criteria were subjected to FLIM imaging. First, imaging was performed using 560 nm laser excitation (30 μW, 40 MHz) for 5 min. Next, a 30-s acquisition was conducted using the 660 nm laser (30 μW, 40 MHz), followed by Acceptor photobleaching using the same laser at a higher power (300 μW, 40 MHz) for 2 min. Post bleaching, the cell was again imaged for 5 min under 560 nm excitation. The same procedure was applied to measure five FG-domain variants. For the variant A221-S338, we were unable to identify cells containing well-formed condensates, possibly due to low GCE efficiency at this site. As a result, we measured an alternative variant, A221-T353, instead.

Regions of interest (ROIs), corresponding to nuclear condensates, were selected based on intensity and average lifetime per pixel (Figure 5d). Time-resolved donor fluorescence intensity profiles were extracted from ROIs before and after acceptor photobleaching. The total fluorescence decay was computed by combining the parallel and perpendicular polarisation components (*I*_∥_ and *I*_⊥_) using^70^:

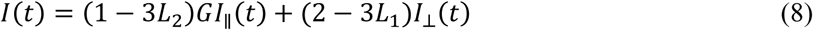

where *L*_1_ and *L*_2_account for polarisation mixing caused by the high-numerical-aperture objective, and *G* is the factor accounting for the difference in the detection efficiencies *η* between parallel and perpendicular polarisation, given by Eq. (9):

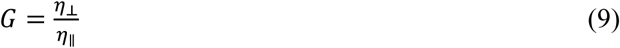

### Lifetime-based analysis of FRET measurements

For cellular measurements, FLIM data were analysed as previously described^40,49^. In brief, the measured fluorescence signal from the donor channels comprises three populations: the donor-only population *I*_Donly_, the FRET population between donor and acceptor dyes *I*_FRET_, and the cellular background signal *I*_bg_. The time-resolved fluorescence intensity in the donor channel can thus be described by Eq. (10).

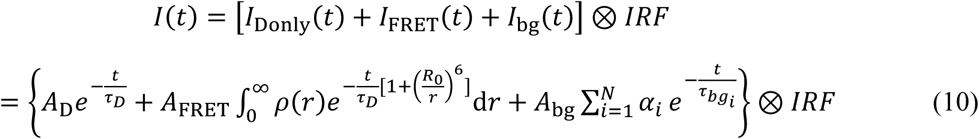

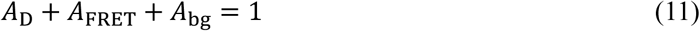

where *A*_D_, *A*_FRET_, and *A*_bg_ represent the amplitude fractions (initial intensities at *t* = 0) of the donor-only, FRET, and background components, respectively. *τ*_*D*_ is the donor lifetime in the absence of FRET, (*r*) describes the inter-residue distance distribution based on a Gaussian chain model, and *R*_0_ is the Förster distance of the dye pair. *τ*_*bgi*_ and *α*_*i*_ characterise the background decay components. The IRF is the instrument response function and was obtained using a freshly prepared saturated solution of potassiumiodide and erythrosine B^71^.

Background lifetimes (*τ*_*bgi*_ and *α*_*i*_) were determined from ∼40 cells expressing the amber-mutant NHA9 in the presence of Boc-lysine, a non-reactive ncAA, and fitted using a four-exponential model via TauFit in PAM^68^. After acceptor photobleaching, FRET contributions are eliminated, simplifying the signal to:

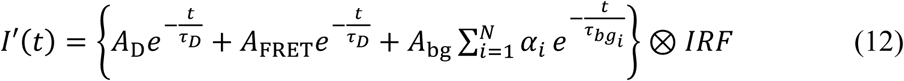

Thus, the background *A*_*bg*_ and *τ*_*D*_ was obtained by fitting with the acceptor photobleaching sample. To isolate *A*_D_, the pre-photobleaching data were fitted using Eq. (10), particularly for samples with short inter-residue distances (i.e., NHA9^A221TAG-S283TAG^ in this case), where FRET signals are clearly distinguishable from the donor-only signals. Following this, the determined *τ*_*D*_ and *A*_bg_, together with 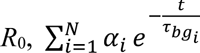, Eq. (10) was used in conjunction with a maximum likelihood estimator (MLE) to extract the root-mean-square inter-residue distance *R*_E_.

Bootstrap resampling was applied to each mutant to account for cellular heterogeneity. In each round, 70% of the data were randomly sampled, and the aggregated fluorescence decay profiles were globally fitted using the Gaussian chain model via MLE in PAM to determine the scaling exponent ν. This resampling and fitting procedure was repeated 10 times. For individual mutant analysis, *R*_E_ was first extracted for each construct, followed by fitting *R*_E_ versus *N*_res_ to derive the apparent scaling exponent ν.

For the lifetime-based analysis of smFRET measurements of nanocluster, background signal was recorded using buffer and *A*_D_ was set to 0 when applying Eq. (10) to extract *R*_E_. In case of lifetime-based analysis of *in vitro* condensates, the background signal was obtained by measuring 10 μM of unlabelled NHA9. To determine *τ*_*D*_ and *A*_D_, 1 nM donor-only labelled NHA9 mixed with 10 μM of unlabelled NHA9 was measured. The extracted *τ*_*D*_ and *A*_D_ was then used to obtain *R*_E_. Fluorescence anisotropy data are presented in Supplementary Figure 9a.

### Phasor plot analysis of FLIM data

The phasor plot, also known as the polar plot, provides a graphical representation of raw FLIM data in a vector space. In this method, each fluorescence decay profile is transformed into a single point on the phasor plot. This transformation is achieved by converting the time-domain fluorescence decay into the frequency domain using a Fourier transform. The phasor coordinates are computed according to the following equations:

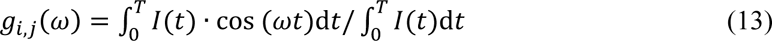

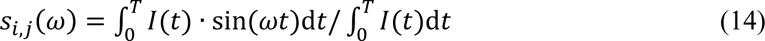

in which *g_i,j_*(ω) and *S_i,j_*(ω) are the *x* and *y* coordinates of the phasor plot, *ω* is the angular frequency of excitation, and *T* is the period of the laser pulses. To ensure accurate scaling of the phasor coordinates, the system was first calibrated by applying a Fourier transform to the measured instrument response function, which was designated as the zero-lifetime reference^72^. Each subsequent phasor point derived from the FLIM data was then calibrated using the same parameters, thereby referencing the final phasor plot to the established calibration standard.

### Molecular Dynamics simulations

The Molecular Dynamics (MD) simulations were carried out using the Hoomd-Blue package version 5.0.1 patched for ROCm 6.3 support^73^. For non-bonded interactions, we used the HPS-Urry force field, which is based on the Urry hydrophobicity scale^55^. Bonded parameters were defined as 3.8 Å and 10kJ/Å² respectively^74^. We used a cutoff radius of 20.0 Å for the ashbaugh-hatch interactions and 35.0 Å for the yukawa potential. The alpha-helices and tertiary structure motif in HOXA9 were kept rigid using elastic networks composed of harmonic bonds with the same bonded parameters^75^. Bonds were created between atoms within less than 12.0 Å from each other. The coordinates for the structured regions are alpha-fold predictions taken from UniProt database (https://www.uniprot.org/uniprotkb/P31269/entry)^76,77^.

All simulations were carried out at 300 K and temperature was kept constant using the Langevin thermostat, with a drag coefficient of 10.0 u/ps. We created cubic boxes with dimensions chosen in order to maintain a fixed volume fraction of 0.01, except for the systems containing 5 chains where a volume fraction of 0.005 was used. The timestep was set to 10 fs.

Individual protein chains were initialised in proximity to each other with random conformations, except for those in the HOXA9 block, which were initialised with the coordinates extracted from the PDB. We then performed an equilibration step consisting of a 100 ns interval where non-bonded interactions are progressively turned on and the timestep was reduced to 0.01 fs and temperature to 150 K, followed by a 24 ns interval where temperature was gradually increased to 300 K, followed by a 56 ns interval where the time step was gradually increased to 10 fs. Throughout all these steps a spherical wall was used to confine the protein chains in order to induce phase separation quicker. We then removed the walls and ran the simulations for 820 ns to further relax the system.

Single chain simulations were set up in a different manner by placing 64 copies in a lattice grid at very large distances from each other in a very large cubic box of side 2400 Å. To ensure that no interaction occurred between the individual chains we fixed the position of the first bead in each chain in space. By following this setup we were able to generate 64 independent single chain trajectories using less computational resources than we would have otherwise used running 64 independent single chain simulations. The equilibration step for single chain simulations was identical to the multi chain one except that no spherical walls were used.

Our production run had a length of 7 μs, on which we carried out our analysis. NHA9 showed a remarkably high phase separation propensity in our simulations, no chains were observed in the dilute phase in any of the simulations. This was confirmed by using a clustering algorithm based on amino acid positions where 2 chains containing at least one pair of amino acids within a distance of less than 7.0 Å were considered to be in the same cluster. The apparent Flory exponent (ν) was computed by fitting the Eq. 6 to the data in Figure 6a.

To compute the density profiles in Fig. 6d and Supplementary Fig. 11, 12, 13 and 14, we first computed the postion of the center of mass (COM) of the cluster at each frame. We then computed histograms of the distance from the COM for each of the 524 amino acids in the sequence. In order to compute the density profiles of each individual block we summed the histograms corresponding to the amino acids in the block, then divided by the number of constituent amino acids of that block to obtain a density in the unit of moles of blocks per liter. To compute contact maps in Supplementary Fig. 15 we considered amino acids to be neighbours if they were within a distance of 1.5σ from each other, where sigma was taken as the mean sigma of the amino acid pair in the ashbaugh-hatch term of the force field, σ = 1/2(σ_1_+σ_2_). We computed the overall interchain contacts in the cluster and then divided them by the number of chains to obtain units of probability of contact per chain.

## Acknowledgements

We acknowledges funding by the Deutsche Forschungsgemeinschaft (DFG, German Research Foundation)–SFB1551–Project No. 464588647, as well as its internal service project on protein production at IMB protein production core facility, especially Martin Möckel and protein analytics at MPI core facility, especially Christine Rosenauer and Svenja Morsbach. E.A.L. also acknowledges funding from the ERC-ADG grant ‘MultiOrganelleDesign’.

## Author contributions

H.R. M.G. S.W. and E.A.L. designed the conceptual framework of the study. H.R. performed the experiments. R.D. conducted molecular dynamics simulations. H.R., R.D., S.W., M.G. and E.A.L. contributed to data acquisition and interpretation. H.R., R.D., S.W., M.G. and E.A.L. wrote the manuscript.

---

**Supplementary Figure 1.**
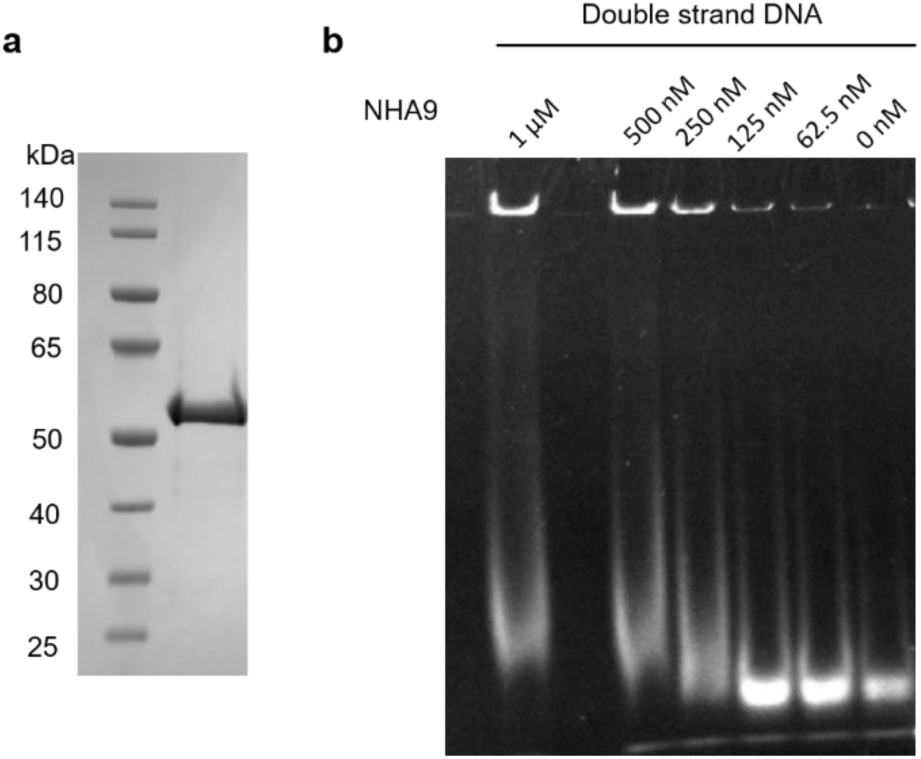
Characterization of purified NHA9. **a** SDS-PAGE analysis of purified NHA9 alongside molecular mass markers. **b** Gel mobility shift assay demonstrating interactions between NHA9 and 20-base pair double-stranded and single-stranded DNA oligonucleotides.

**Supplementary Figure 2.**
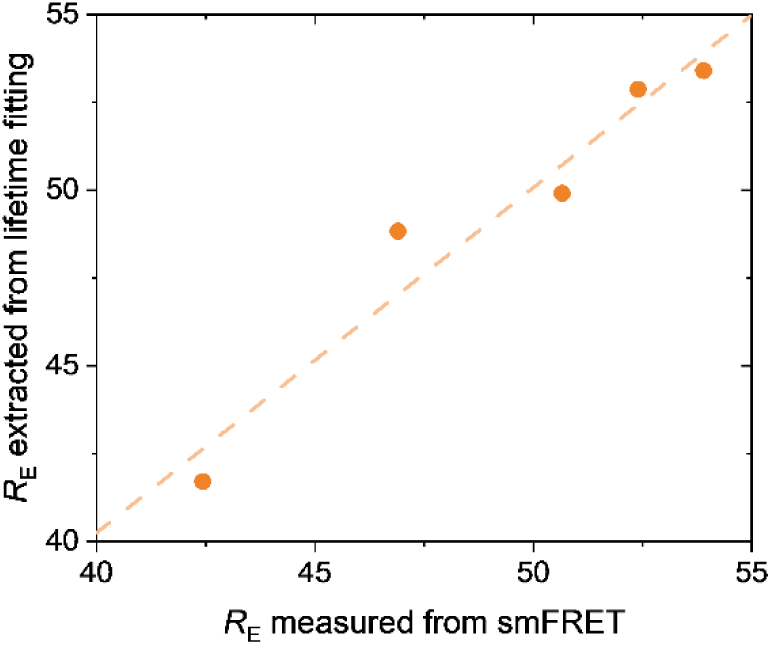
Correlation between inter-residue distance *R*_E_ measured by intensity-based single-molecule FRET and those obtained from lifetime-based fitting. The red dotted line indicates the fitted correlation curve. The high correlation coefficient (*R*^2^ = 0.95) validates the accuracy and reliability of our FLIM-FRET pipeline for measuring *R*_E_.

**Supplementary Figure 3.**
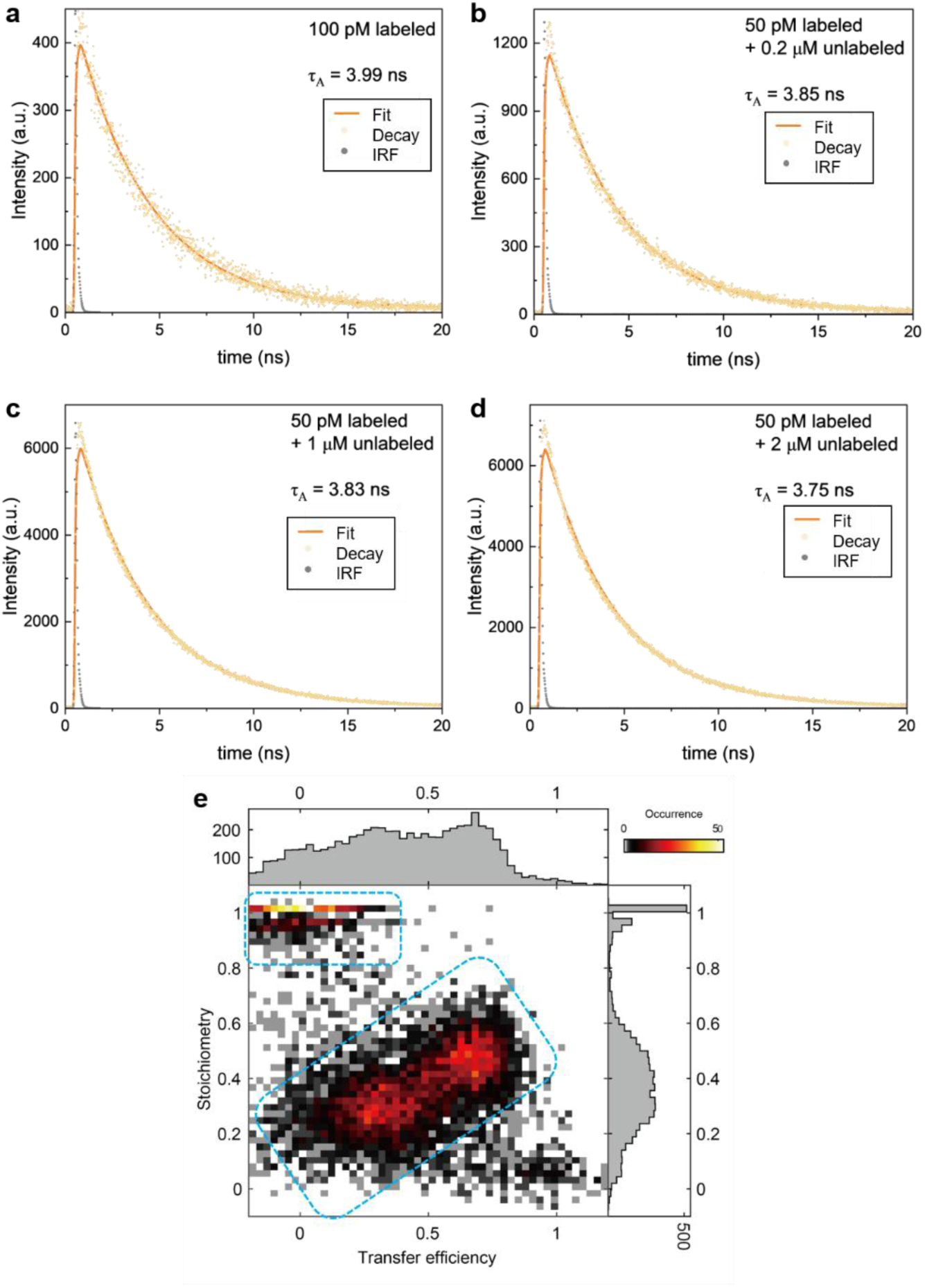
Fluorescence lifetime decays and fitting of Alexa 594 acceptor conjugated to NHA9 at different concentrations and a 2D histogram of stoichiometry versus transfer efficiency values. Lifetime decays were obtained from single-molecule FRET measurements during the acceptor excitation cycle. **a** The fluorescence lifetime of Alexa 594 conjugated to NHA9 in the monomeric state was 3.99 ns. Increasing NHA9 concentrations resulted in decreased lifetimes: **b** 3.85 ns at 0.2 µM, **c** 3.83 ns at 1 µM, and **d** 3.75 ns at 2 µM. These data indicate that the acceptor undergoes fluorescence quenching within nanoclusters. **e** The 2D histogram of the stoichiometry values versus the transfer efficiency for NHA9_A221-S362_ at 1 µM concentration. The Donor-only species and FRET species, highlighted by blue rectangles, were individually selected for separate fitting using a 2D Gaussian function.

**Supplementary Figure 4.**
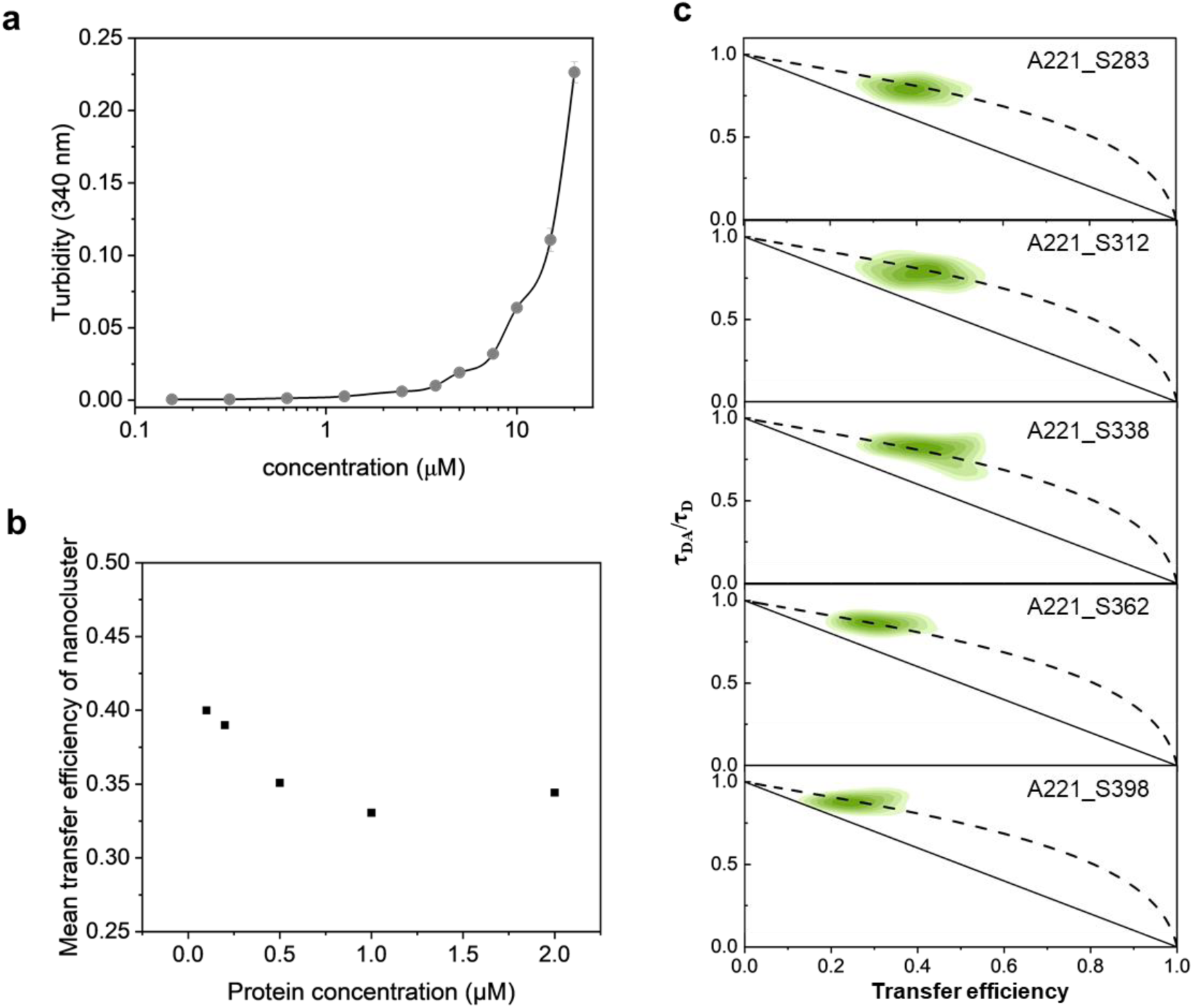
Turbidity assay and nanocluster analysis by smFRET. **a** concentration-dependent turbidity of purified NHA9, measured by UV absorbance at 340 nm. **b** Mean transfer efficiency of NHA9 nanocluster at varying concentrations. **c** Two-dimensional histogram of the relative donor fluorescence lifetime (τ_DA_/τ_D_) versus transfer efficiency (*E*) calculated from individual bursts of five nanocluster variants at 1 µM. The dashed line indicates the dynamic relationship predicted by a Gaussian chain polymer model. See Methods for additional details.

**Supplementary Figure 5.**
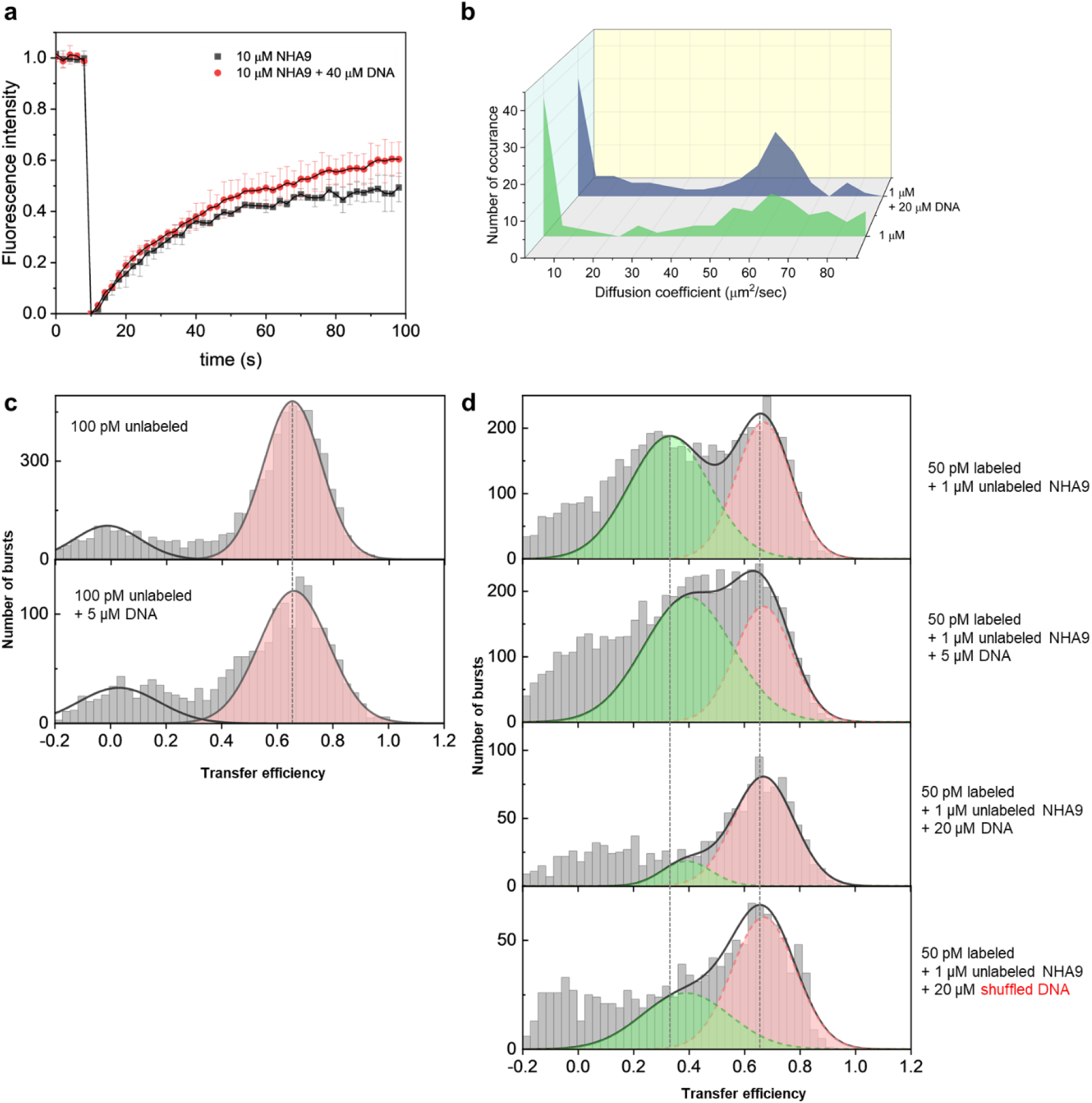
Effects of DNA fragments on NHA9 condensate and nanocluster formation. **a** FRAP recovery curves of NHA9 condensates in the presence and absence of DNA fragment. Data are presented as mean ± SD (*n* = 3). **b** Nanocluster formation of NHA9 with or without DNA fragment at 1 µM, assessed by FCS. Autocorrelation curves were derived from 120 repeated measurements with short sampling time (10 s), using NHA9_A221C_ labeled with LD655 as fluorescent probe in trace amount (10 nM) with excess unlabeled NHA9. **c** Single-molecule transfer efficiency histograms of NHA9_A221-S362_ at 100 pM with or without 5 µM DNA fragments. **d** Single-molecule transfer efficiency histograms of NHA9_A221-S362_ at 1 µM unlabeled protein with varying concentrations of DNA fragments. The shuffled DNA fragment (5’-CTATATGCTATCGTGTAACC-3’) only partially inhibited NHA9 nanocluster formation, suggesting the coexistence of specific and non-specific interactions between NHA9 and the double-stranded recognition DNA fragment.

**Supplementary Figure 6.**
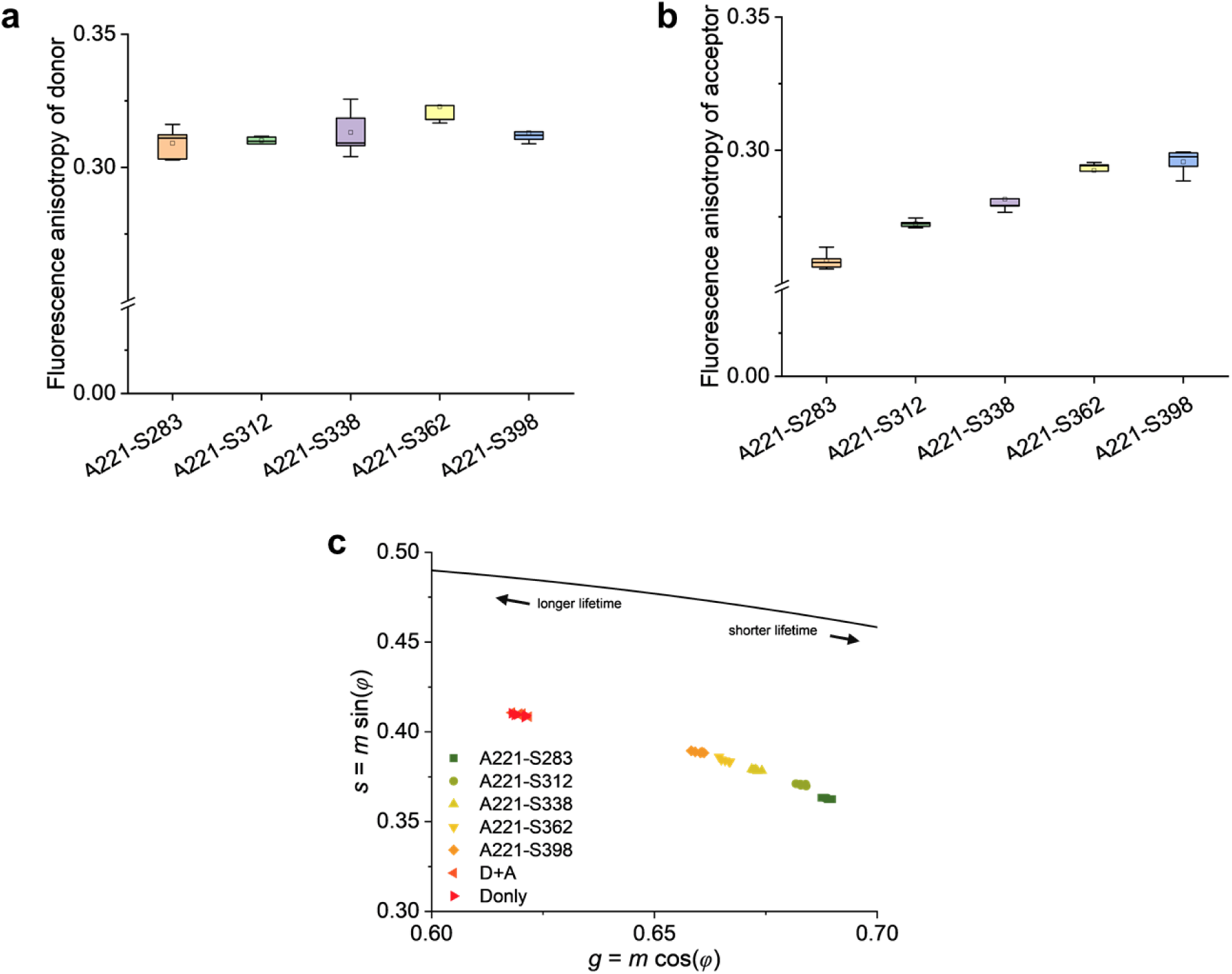
FLIM-FRET analysis of NHA9 condensates *in vitro*. NHA9 variants were site-specifically labelled with a FRET dye pair, with 1 nM labeled protein mixed with 10 µM unlabelled protein. Each variant was analyzed in five replicates. **a** Box plot of fluorescence anisotropy values of the donor dye at different labeling sites in NHA9 condensates. **b** Box plot of fluorescence anisotropy values of the acceptor dye at different labeling sites in NHA9 condensates. Donor dye anisotropy values exceeded 0.3 across all sites, while acceptor dye anisotropy values remained below 0.3. In this regime, the error in distance measurements is estimated to be less than 10%^1^. **c** Phasor plots of donor fluorescence signals from NHA9 variants in *in vitro* reconstituted condensates. Each sample was measured for 5 repeats. No differences were observed between donor-only and donor–acceptor labeled single mutants, indicating the absence of intermolecular FRET events.

**Supplementary Figure 7.**
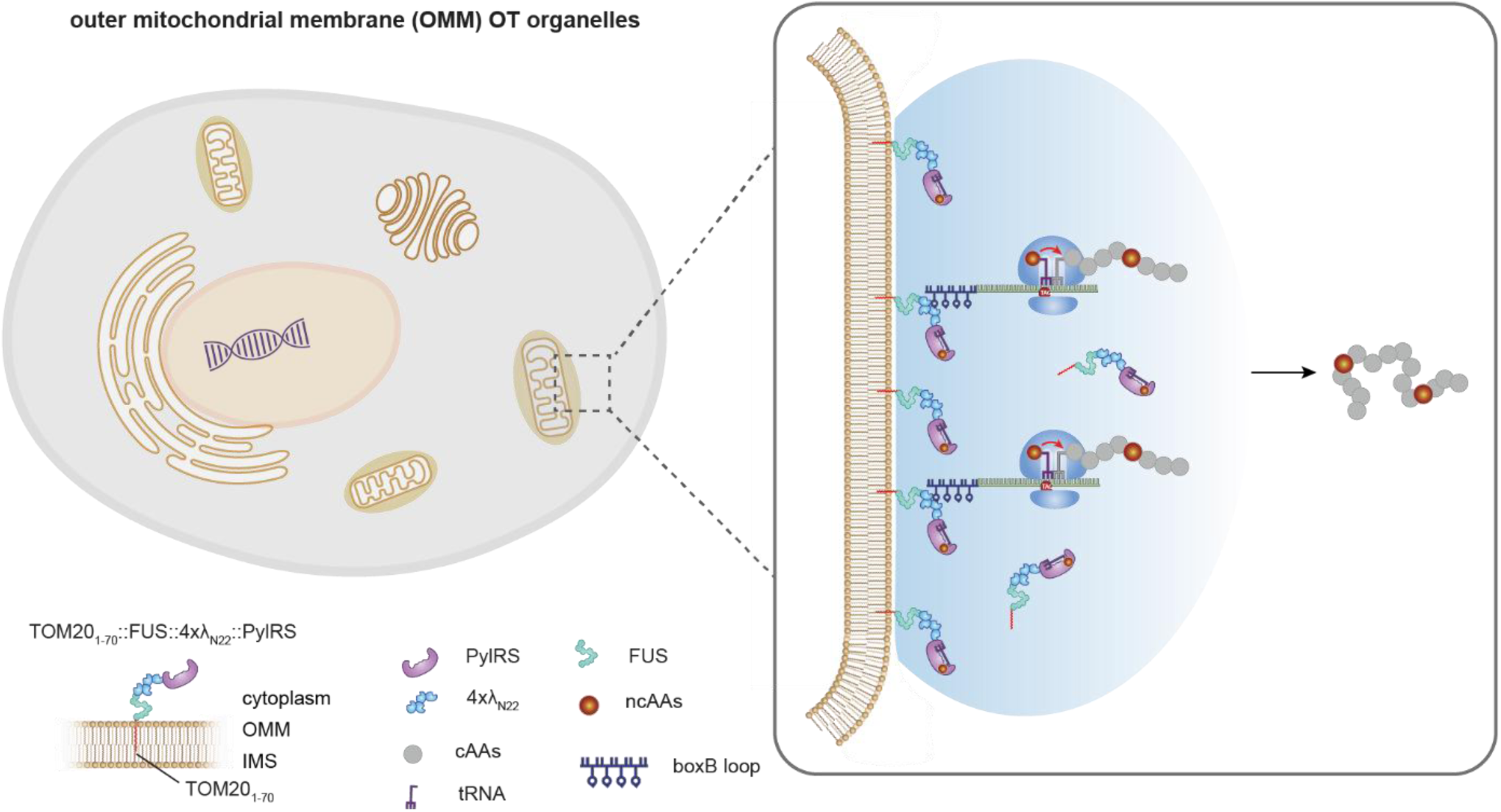
Schematic of NHA9 labeling by outer mitochondrial membrane OTO-GCE system. PylRS and λ_N22_ were fused to the phase-separating scaffold FUS and the mitochondrial outer membrane-targeting domain TOM20_1-70_. NHA9 mRNA was engineered with boxB RNA hairpin loops, which specifically bind to the λ_N22_ peptides. This design enabled selective recruitment of NHA9 mRNA into OTO system, thereby restricting the incorporation of ncAAs to NHA9 and preventing their incorporation into other proteins.

**Supplementary Figure 8.**
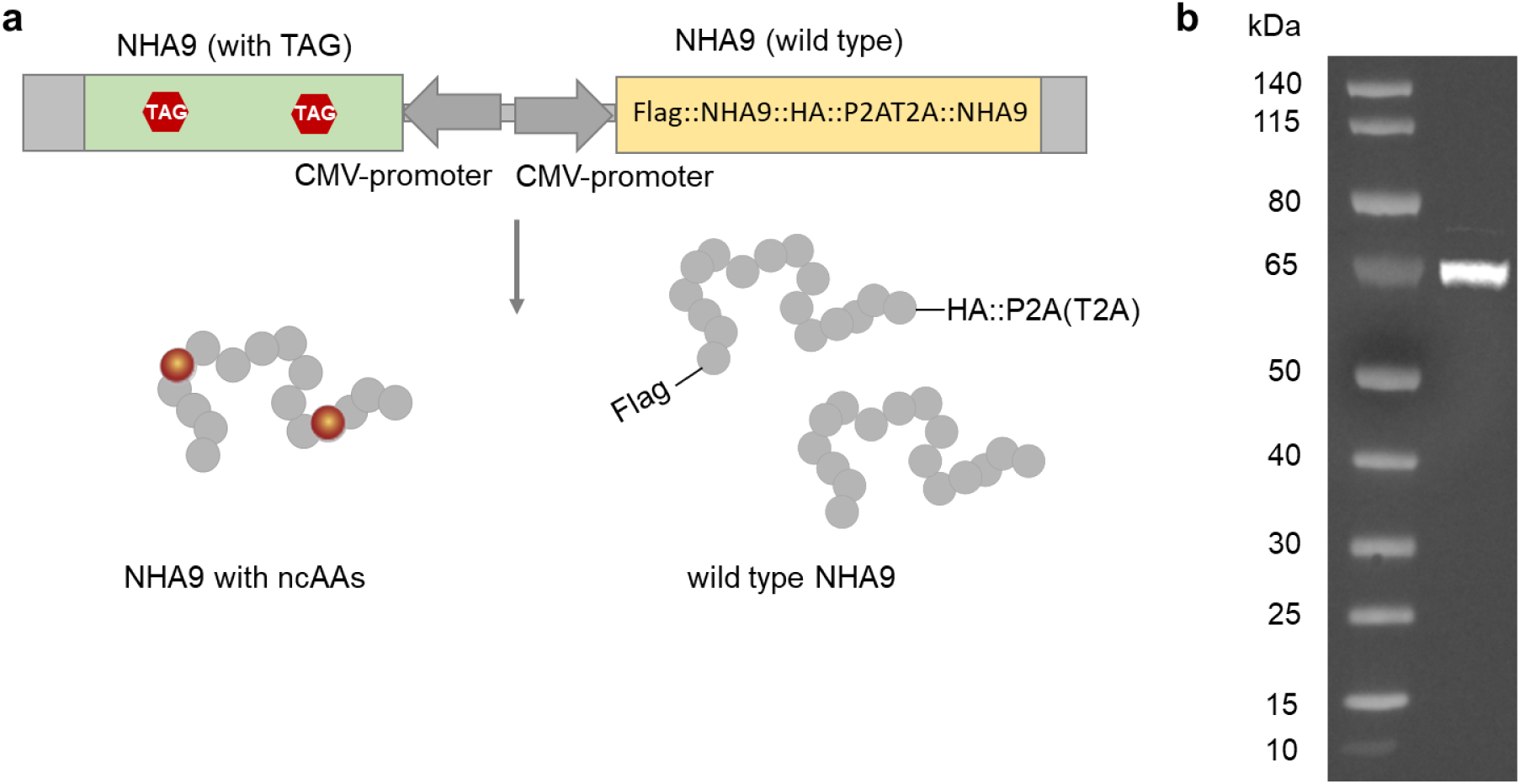
NHA9 expression in HeLa cells. **a** Schematic representation of constructs used for NHA9 expression. To account for the heterogeneity of transient transfection, the construct was designed to co-express both wild-type NHA9 and TAG-mutant NHA9 within the same cells. **b** Western blot analysis of wild-type NHA9 expression using an anti-HA antibody. Due to ribosomal skipping induced by 2A peptides, cells expressed either Flag-NHA9-HA-P2A-T2A (65.8 kDa) or Flag-NHA9-HA-P2A (63.8 kDa), depending on the specific site of skipping.

**Supplementary Figure 9.**
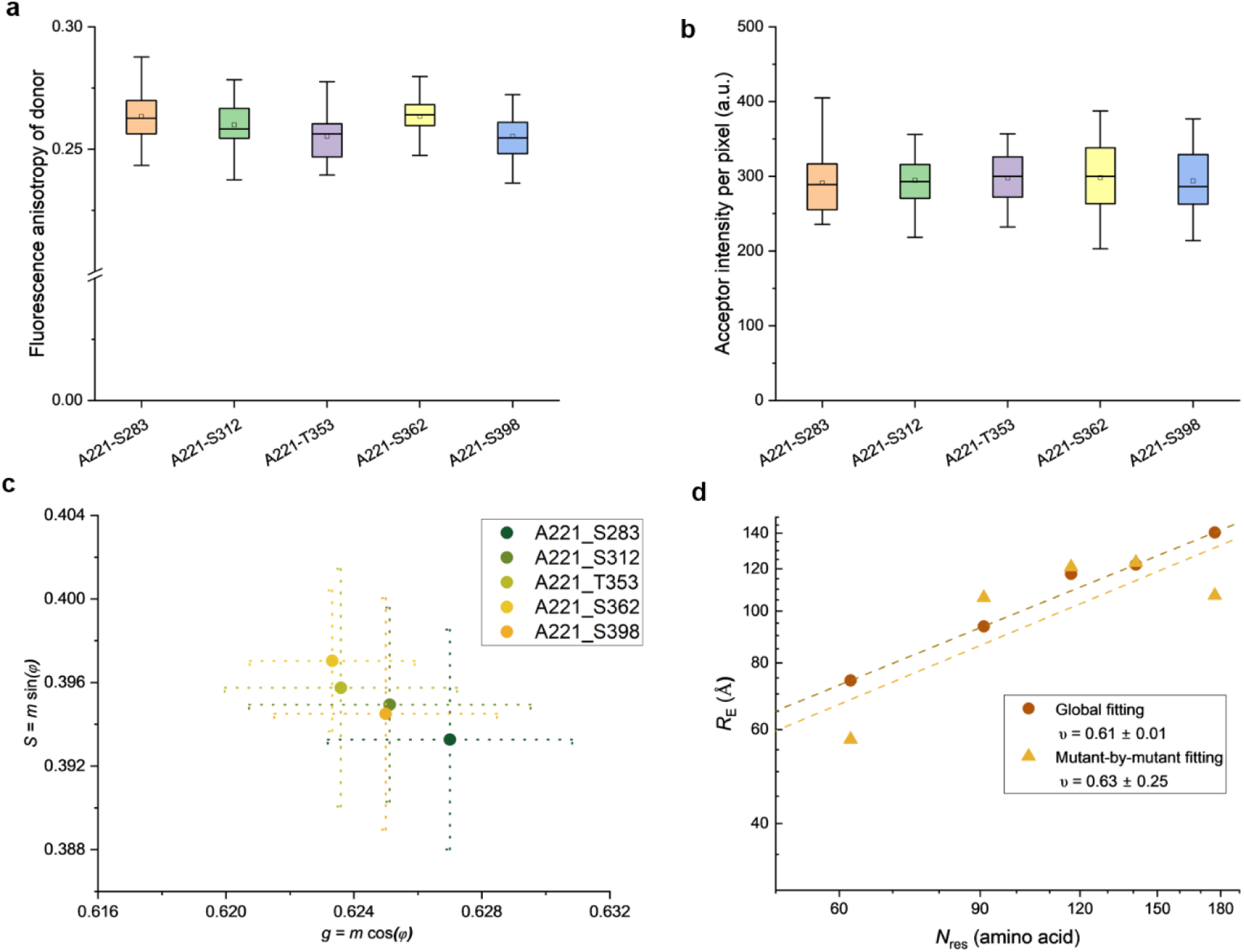
Analysis of FLIM-FRET data in cells. **a** Box plot of donor anisotropy measurements across different NHA9 variants. All anisotropy values were below 0.3, indicating sufficient rotational mobility for FRET measurements of the donor fluorophore in the cellular environment. **b** Box plot of acceptor fluorescence intensity per pixel within nuclear condensates under 660-nm laser excitation. All five variants exhibited similar expression levels, with no cells displaying abnormally high expression included in the dataset. **c** Phasor plots of fluorescence donor channel signals for NHA9 variants. **d** Comparison of the *R*_E_ obtained from global fitting versus mutant-by-mutant fitting using the Gaussian chain model. In global fitting, fluorescence lifetime decays from all variants were simultaneously fitting using Eq. 6 to extract the scaling exponent across all measured cells. In mutant-by-mutant fitting, *R*_E_ values were first individual extracted for each variant by fitting its respective lifetime decay using Eq. 9, and the scaling exponent was then determined by fitting extracting *R*_E_ versus *N*_res_ using Eq. 6.

**Supplementary Figure 10.**
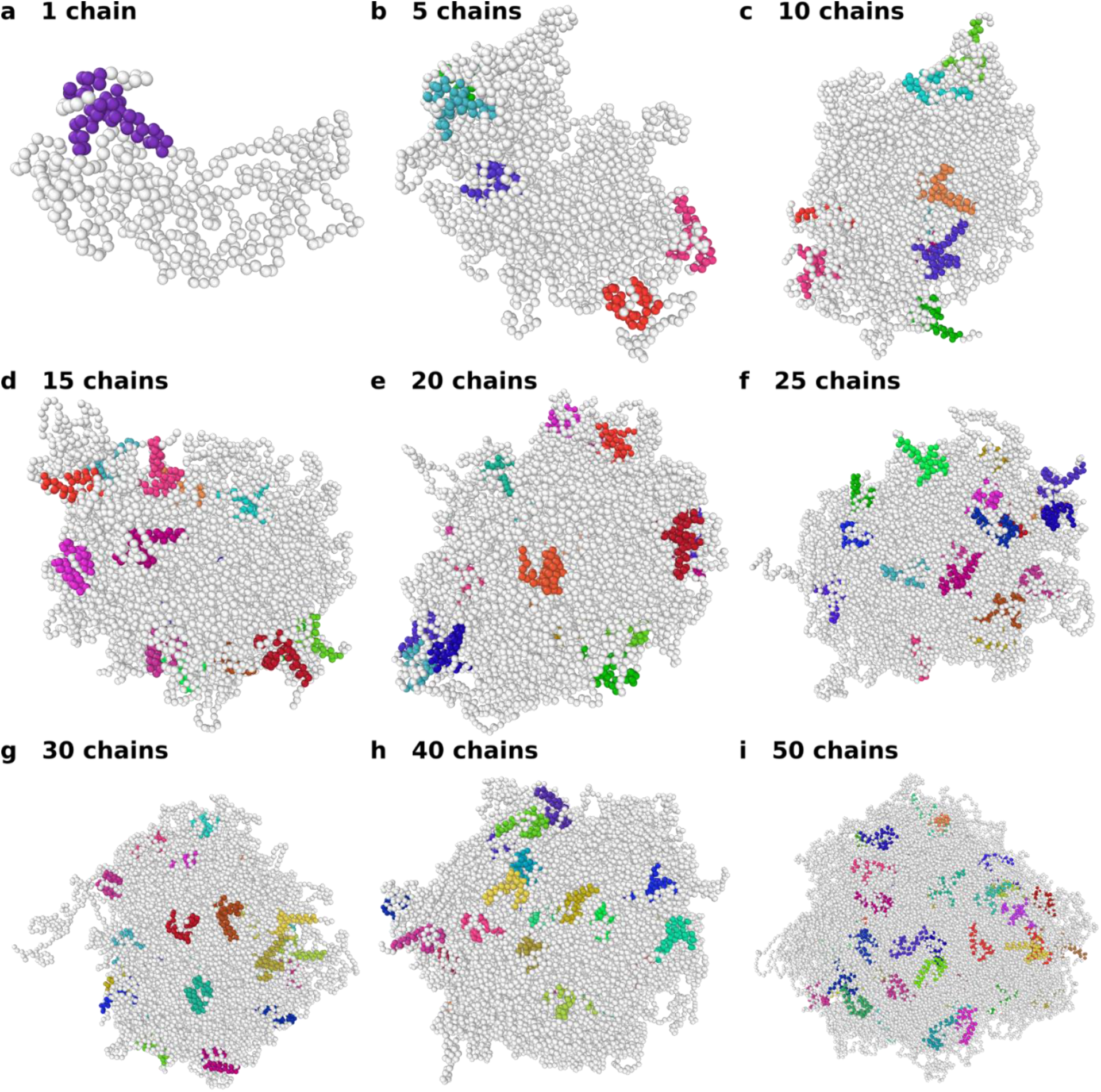
Snapshots of clusters of different sizes. The 3 alpha-helices making up the DBD are colored, different colors represent different protein chains.

**Supplementary Figure 11.**
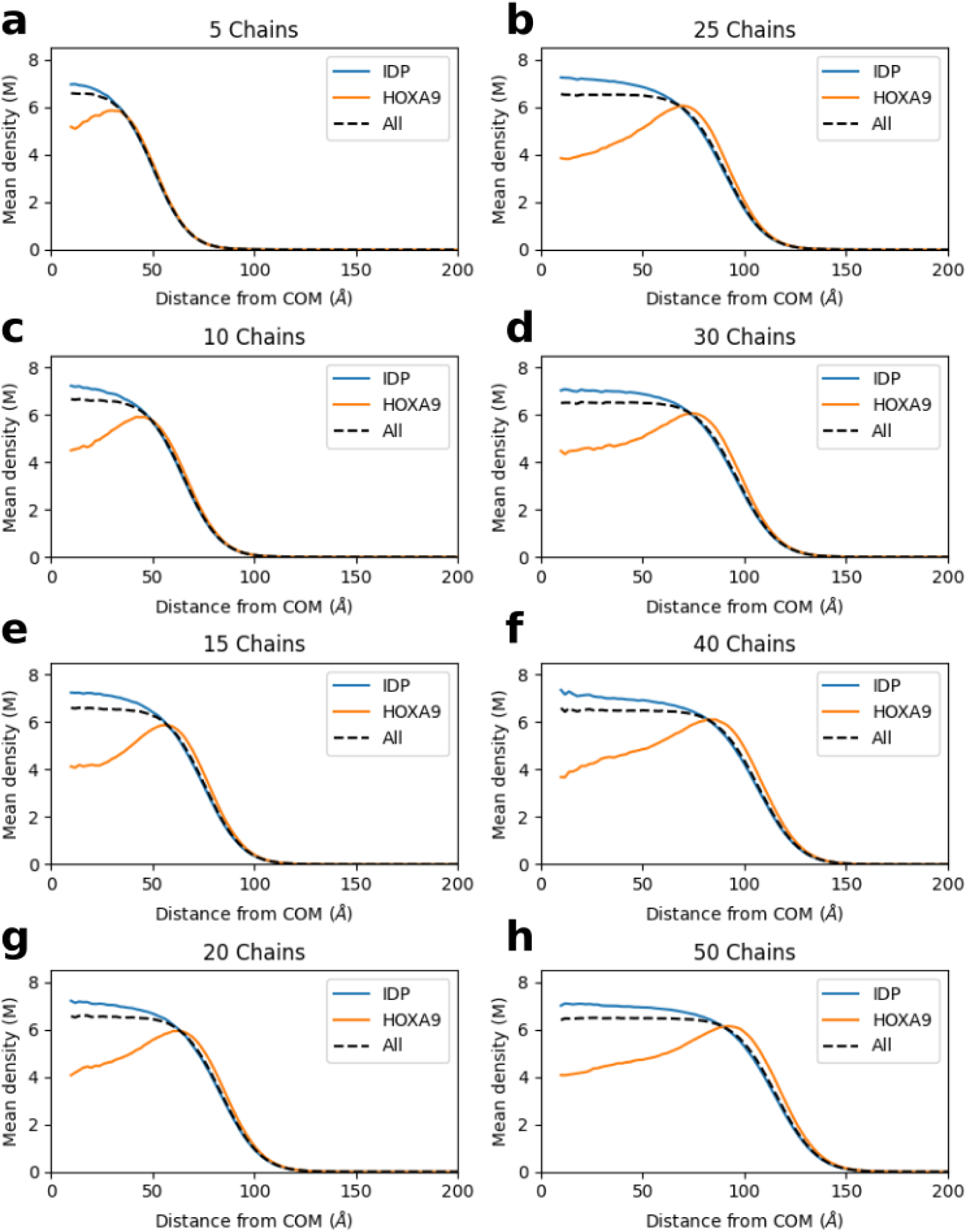
Density profiles for individual clusters. Blue lines represent the IDP, orange lines represent the HOXA9 block and the dashed black lines correspond to full-length NHA9.

**Supplementary Figure 12.**
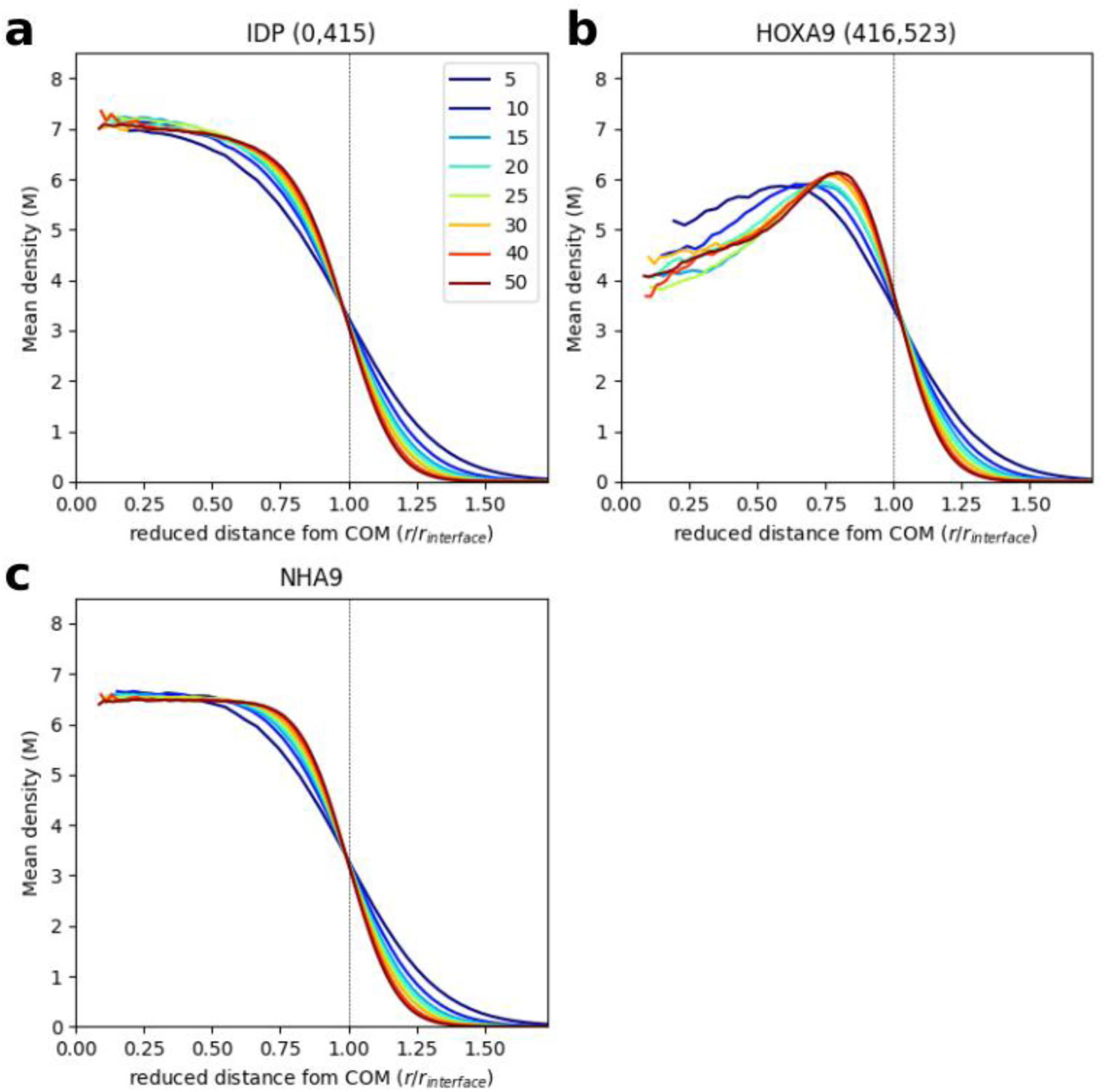
Density profiles for both blocks of NHA9. **a** IDP block, **b** HOXA9 block and **c** full protein. Colors indicate clusters of different sizes. The dashed black line represents the center of the cluster’s interface. The numbers next to the block names indicate the indices of the first and last amino acid in that block.

**Supplementary Figure 13.**
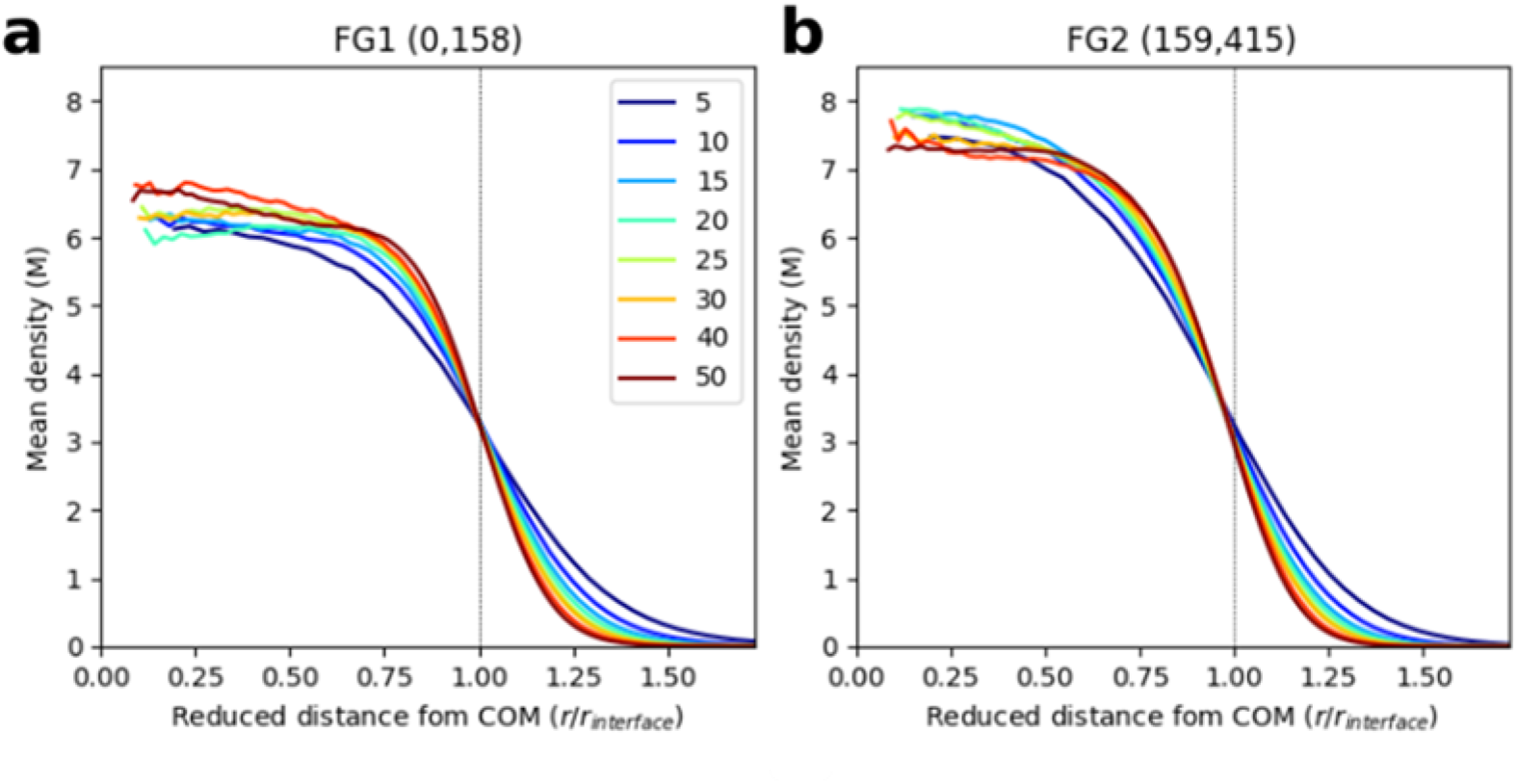
Density profiles with the IDP divided into two blocks. **a** FG1 block, **b** FG2 block. The FG1 region includes the chain end of the IDP. Colors indicate clusters of different sizes. The dashed black line represents the center of the cluster’s interface. The numbers next to the block names indicate the indices of the first and last amino acid in that block.

**Supplementary Figure 14.**
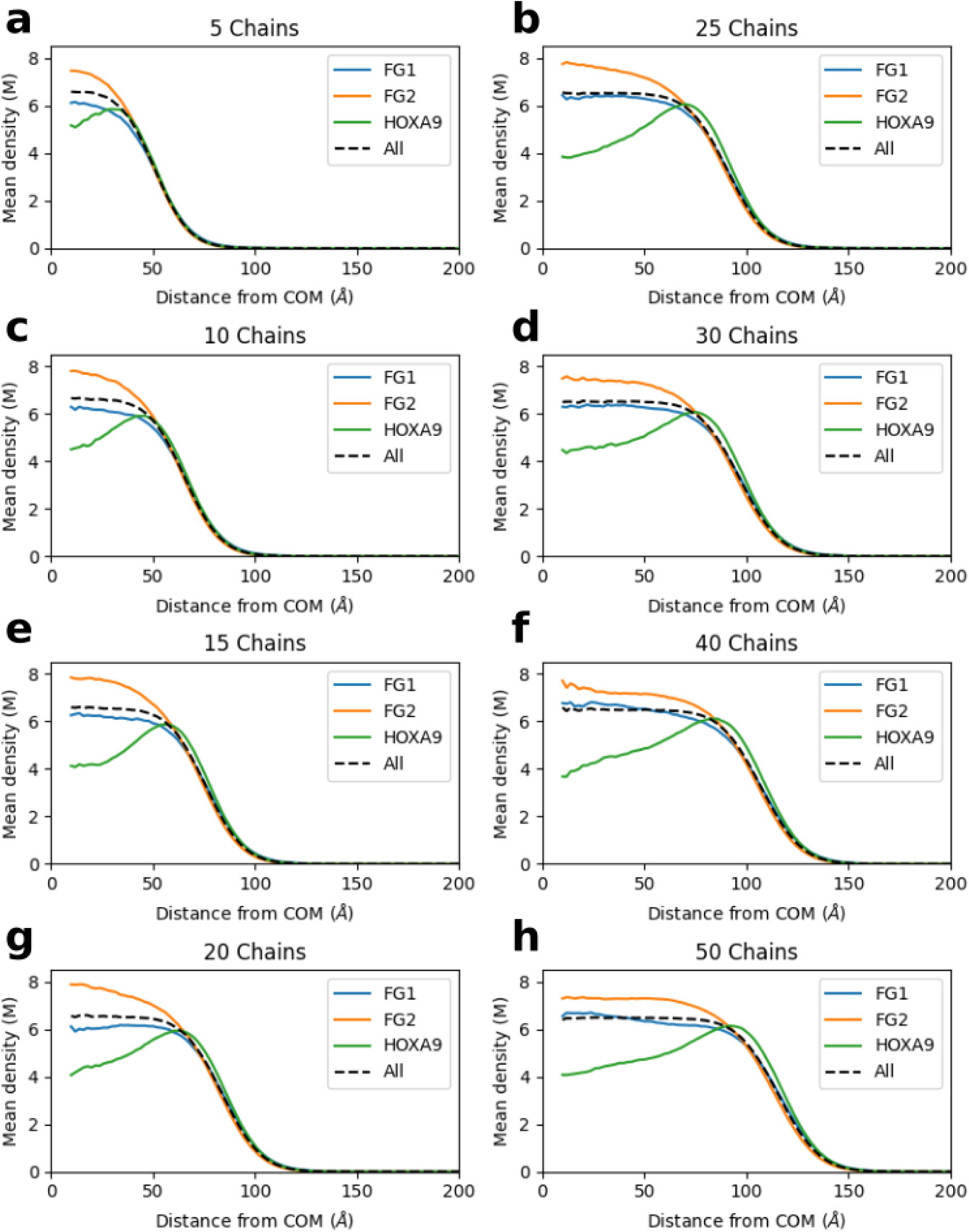
Density profiles of individual clusters. Blue and orange lines represent the FG1 and FG2 regions of the IDP block, respectively. Green lines represent HOXA9, and the dashed black lines correspond to full-length NHA9. FG1 is located at the chain end of the IDP block.

**Supplementary Figure 15.**
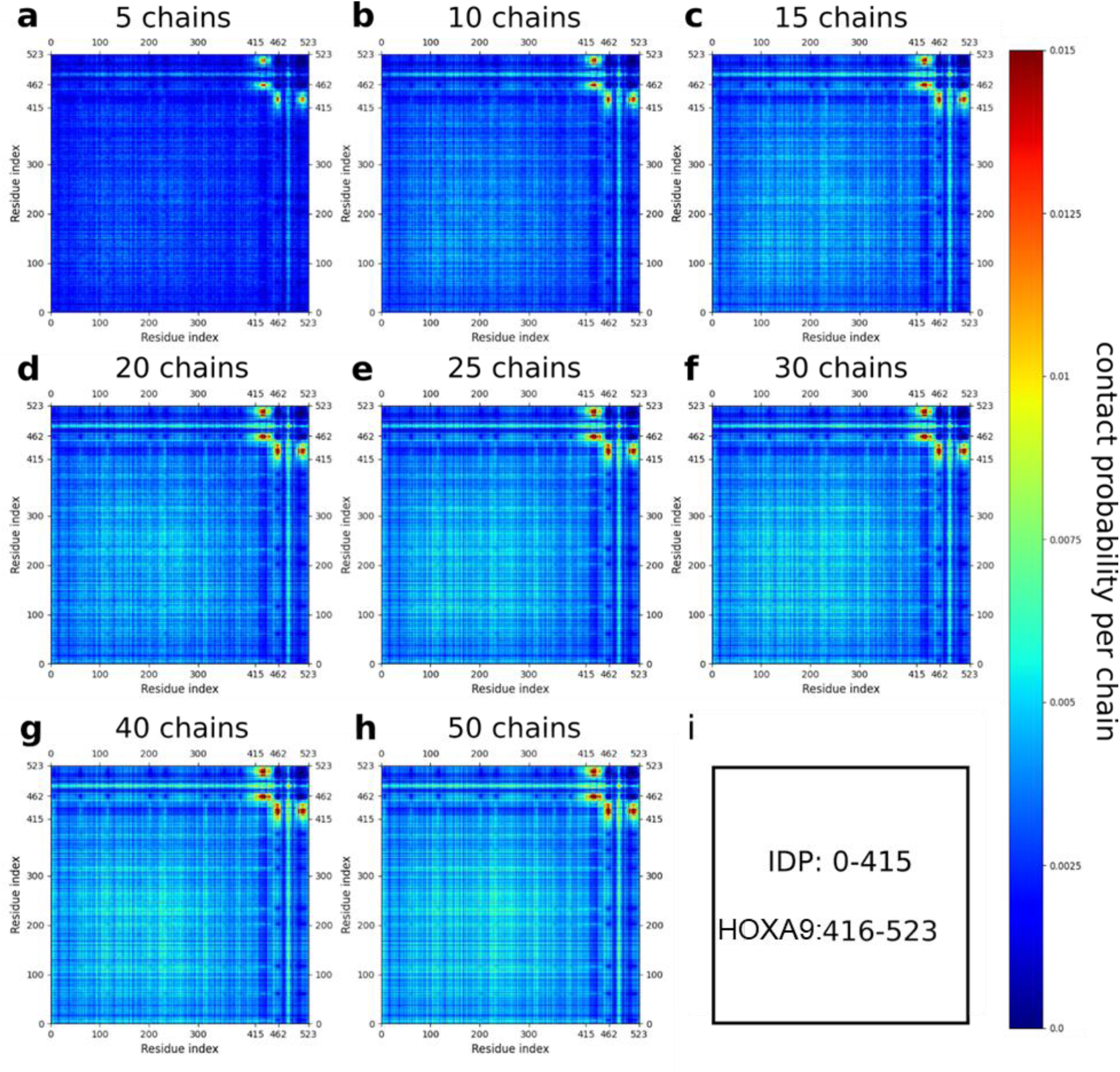
Interchain contact map for clusters of different sizes. **a** 5, **b** 10, **c** 15, **d** 20, **e** 25, **f** 30, **g** 40, **h** 50 chains. **i** Indices of the amino acids making up each of the blocks/regions of NHA9

## References

1. Albert, B., et al. Molecular Biology of the Cell. (Garland Science).

2. Lambert, S. A. et al. The Human Transcription Factors. Cell 172, 650–665 (2018).

3. Ferrie, J. J., Karr, J. P., Tjian, R. & Darzacq, X. “Structure”-function relationships in eukaryotic transcription factors: The role of intrinsically disordered regions in gene regulation. Mol. Cell 82, 3970–3984 (2022).

4. Liu, J. et al. Intrinsic Disorder in Transcription Factors. Biochem. 45, 6873–6888 (2006).

5. Xhani, S. et al. Intrinsic disorder controls two functionally distinct dimers of the master transcription factor PU.1. Sci. Adv. 6, eaay3178 (2020).

6. Lobley, A., Swindells, M. B., Orengo, C. A. & Jones, D. T. Inferring Function Using Patterns of Native Disorder in Proteins. PLoS Comput. Biol. 3, e162 (2007).

7. Pei, G., Lyons, H., Li, P. & Sabari, B. R. Transcription regulation by biomolecular condensates. Nat. Rev. Mol. Cell Biol. (2024) doi:10.1038/s41580-024-00789-x.

8. Boija, A. et al. Transcription Factors Activate Genes through the Phase-Separation Capacity of Their Activation Domains. Cell 175, 1842–1855.e16 (2018).

9. Cho, W.-K. et al. Mediator and RNA polymerase II clusters associate in transcription-dependent condensates. Science 361, 412–415 (2018).

10. Chong, S. et al. Imaging dynamic and selective low-complexity domain interactions that control gene transcription. Science 361, eaar2555 (2018).

11. Sabari, B. R. et al. Coactivator condensation at super-enhancers links phase separation and gene control. Science 361, eaar3958 (2018).

12. Wei, M.-T. et al. Nucleated transcriptional condensates amplify gene expression. Nat. Cell Biol. 22, 1187–1196 (2020).

13. Ma, L. et al. Co-condensation between transcription factor and coactivator p300 modulates transcriptional bursting kinetics. Mol. Cell 81, 1682–1697.e7 (2021).

14. Du, M. et al. Direct observation of a condensate effect on super-enhancer controlled gene bursting. Cell 187, 331–344.e17 (2024).

15. Trojanowski, J. et al. Transcription activation is enhanced by multivalent interactions independent of phase separation. Mol. Cell 82, 1878–1893.e10 (2022).

16. He, J. et al. Dual-role transcription factors stabilize intermediate expression levels. Cell 187, 2746–2766.e25 (2024).

17. Zuo, L. et al. Loci-specific phase separation of FET fusion oncoproteins promotes gene transcription. Nat. Commun. 12, 1491 (2021).

18. Zhang, H. et al. Reversible phase separation of HSF1 is required for an acute transcriptional response during heat shock. Nat. Cell Biol. 24, 340–352 (2022).

19. Hu, X. et al. Nuclear condensates of YAP fusion proteins alter transcription to drive ependymoma tumourigenesis. Nat. Cell Biol. (2023) doi:10.1038/s41556-022-01069-6.

20. Sanborn, A. L. et al. Simple biochemical features underlie transcriptional activation domain diversity and dynamic, fuzzy binding to Mediator. eLife 10, e68068 (2021).

21. Udupa, A., Kotha, S. R. & Staller, M. V. Commonly asked questions about transcriptional activation domains. Curr. Opin. Struct. Biol. 84, 102732 (2024).

22. Perry, J. A., Seong, B. K. A. & Stegmaier, K. Biology and Therapy of Dominant Fusion Oncoproteins Involving Transcription Factor and Chromatin Regulators in Sarcomas. Annu. Rev. Cancer Biol. 3, 299–321 (2019).

23. Gough, S. M., Slape, C. I. & Aplan, P. D. NUP98 gene fusions and hematopoietic malignancies: common themes and new biologic insights. Blood 118, 6247–6257 (2011).

24. Nakamura, T. et al. Fusion of the nucleoporin gene NUP98 to HOXA9 by the chromosome translocation t(7;11)(p15;p15) in human myeloid leukaemia. Nat. Genet. 12, 154–158 (1996).

25. Yamamoto, K., Nakamura, Y., Nakamura, Y., Saito, K. & Furusawa, S. Expression of the NUP98/HOXA9 fusion transcript in the blast crisis of Philadelphia chromosome-positive chronic myelogenous leukaemia with t(7;11)(p15;p15). Br. J. Haematol. 109, 423–426 (2000).

26. Kroon, E. NUP98-HOXA9 expression in hemopoietic stem cells induces chronic and acute myeloid leukemias in mice. EMBO J. 20, 350–361 (2001).

27. Radu, A., Moore, M. S. & Blobel, G. The peptide repeat domain of nucleoporin Nup98 functions as a docking site in transport across the nuclear pore complex. Cell 81, 215–222 (1995).

28. Terlecki-Zaniewicz, S. et al. Biomolecular condensation of NUP98 fusion proteins drives leukemogenic gene expression. Nat. Struct. Mol. Biol. 28, 190–201 (2021).

29. Ahn, J. H. et al. Phase separation drives aberrant chromatin looping and cancer development. Nature 595, 591–595 (2021).

30. Chandra, B. et al. Phase Separation Mediates NUP98 Fusion Oncoprotein Leukemic Transformation. Cancer Discov. 12, 1152–1169 (2022).

31. Oka, M. et al. Phase-separated nuclear bodies of nucleoporin fusions promote condensation of MLL1/CRM1 and rearrangement of 3D genome structure. Cell Rep. 42, 112884 (2023).

32. Kar, M. et al. Phase-separating RNA-binding proteins form heterogeneous distributions of clusters in subsaturated solutions. Proc. Natl. Acad. Sci. U.S.A. 119, e2202222119 (2022).

33. Ray, S. et al. Mass photometric detection and quantification of nanoscale α-synuclein phase separation. Nat. Chem. 15, 1306–1316 (2023).

34. Lan, C. et al. Quantitative real-time in-cell imaging reveals heterogeneous clusters of proteins prior to condensation. Nat. Commun. 14, 4831 (2023).

35. Wollman, A. J. et al. Transcription factor clusters regulate genes in eukaryotic cells. eLife 6, e27451 (2017).

36. Lerner, E. et al. Toward dynamic structural biology: Two decades of single-molecule Förster resonance energy transfer. Science 359, eaan1133 (2018).

37. Hellenkamp, B. et al. Precision and accuracy of single-molecule FRET measurements—a multi-laboratory benchmark study. Nat. Methods 15, 669–676 (2018).

38. Agam, G. et al. Reliability and accuracy of single-molecule FRET studies for characterization of structural dynamics and distances in proteins. Nat. Methods 20, 523–535 (2023).

39. Ng, S. C., Güttler, T. & Görlich, D. Recapitulation of selective nuclear import and export with a perfectly repeated 12mer GLFG peptide. Nat. Commun. 12, 4047 (2021).

40. Yu, M. et al. Visualizing the disordered nuclear transport machinery in situ. Nature 617, 162–169 (2023).

41. Hoffmann, A. et al. Mapping protein collapse with single-molecule fluorescence and kinetic synchrotron radiation circular dichroism spectroscopy. Proc. Natl. Acad. Sci. U.S.A. 104, 105–110 (2007).

42. Fuertes, G. et al. Decoupling of size and shape fluctuations in heteropolymeric sequences reconciles discrepancies in SAXS vs. FRET measurements. Proc. Natl. Acad. Sci. U.S.A. 114, (2017).

43. Flory, P. J. The Configuration of Real Polymer Chains. J. Chem. Phys. 17, 303–310 (1949).

44. Ribbeck, K. The permeability barrier of nuclear pore complexes appears to operate via hydrophobic exclusion. EMBO J. 21, 2664–2671 (2002).

45. Le Ferrand, H., Duchamp, M., Gabryelczyk, B., Cai, H. & Miserez, A. Time-Resolved Observations of Liquid–Liquid Phase Separation at the Nanoscale Using *in Situ* Liquid Transmission Electron Microscopy. J. Am. Chem. Soc. 141, 7202–7210 (2019).

46. Pittman, J. M. et al. Nanodroplet Oligomers (NanDOs) of Aβ40. Biochem. 60, 2691–2703 (2021).

47. Zhao, H. et al. Energetic and structural features of SARS-CoV-2 N-protein co-assemblies with nucleic acids. iScience 24, 102523 (2021).

48. Peng, S. et al. Phase separation at the nanoscale quantified by dcFCCS. Proc. Natl. Acad. Sci. U.S.A. 117, 27124–27131 (2020).

49. Ruan, H. & Lemke, E. A. Resolving Conformational Plasticity in Mammalian Cells with High-Resolution Fluorescence Tools. Annu. Rev. Phys. Chem. 76, 103–128 (2025).

50. Zhou, H.-X., Rivas, G. & Minton, A. P. Macromolecular Crowding and Confinement: Biochemical, Biophysical, and Potential Physiological Consequences. Annu. Rev. Biophys. 37, 375–397 (2008).

51. Plitzko, J. M., Schuler, B. & Selenko, P. Structural Biology outside the box — inside the cell. Curr. Opin. Struct. Biol. 46, 110–121 (2017).

52. Nikić, I. & Lemke, E. A. Genetic code expansion enabled site-specific dual-color protein labeling: superresolution microscopy and beyond. Curr. Opin. Chem. Biol. 28, 164–173 (2015).

53. Reinkemeier, C. D., Girona, G. E. & Lemke, E. A. Designer membraneless organelles enable codon reassignment of selected mRNAs in eukaryotes. Science 363, eaaw2644 (2019).

54. Reinkemeier, C. D. & Lemke, E. A. Dual film-like organelles enable spatial separation of orthogonal eukaryotic translation. Cell 184, 4886–4903.e21 (2021).

55. Regy, R. M., Thompson, J., Kim, Y. C. & Mittal, J. Improved coarse-grained model for studying sequence dependent phase separation of disordered proteins. Protein Sci. 30, 1371–1379 (2021).

56. Wang, J. et al. Sequence-dependent conformational transitions of disordered proteins during condensation. Chem. Sci. 15, 20056–20063 (2024).

57. Wu, T. et al. Single-fluorogen imaging reveals distinct environmental and structural features of biomolecular condensates. Nat. Phys. (2025) doi:10.1038/s41567-025-02827-7.

58. Wang, J., Devarajan, D. S., Nikoubashman, A. & Mittal, J. Conformational Properties of Polymers at Droplet Interfaces as Model Systems for Disordered Proteins. ACS Macro Lett. 12, 1472–1478 (2023).

59. Hnisz, D., Shrinivas, K., Young, R. A., Chakraborty, A. K. & Sharp, P. A. A Phase Separation Model for Transcriptional Control. Cell 169, 13–23 (2017).

60. Mizutani, A., Tan, C., Sugita, Y. & Takada, S. Micelle-like clusters in phase-separated Nanog condensates: A molecular simulation study. PLoS Comput. Biol. 19, e1011321 (2023).

61. Shinn, M. K. et al. Nuclear speckle proteins form intrinsic and MALAT1-dependent microphases. Preprint at 10.1101/2025.02.26.640430 (2025).

62. Ianiro, A. et al. Liquid–liquid phase separation during amphiphilic self-assembly. Nat. Chem. 11, 320–328 (2019).

63. Sato, T. & Takahashi, R. Competition between the micellization and the liquid–liquid phase separation in amphiphilic block copolymer solutions. Polym. J. 49, 273–277 (2017).

64. Chong, S. et al. Tuning levels of low-complexity domain interactions to modulate endogenous oncogenic transcription. Mol. Cell 82, 2084–2097.e5 (2022).

65. Lemke, E. A. Site-Specific Labeling of Proteins for Single-Molecule FRET Measurements Using Genetically Encoded Ketone Functionalities. in Bioconjugation Protocols (ed. Mark, S. S.) vol. 751 3–15 (Humana Press, Totowa, NJ, 2011).

66. Kudryavtsev, V. et al. Combining MFD and PIE for Accurate Single-Pair Förster Resonance Energy Transfer Measurements. ChemPhysChem 13, 1060–1078 (2012).

67. Müller, B. K., Zaychikov, E., Bräuchle, C. & Lamb, D. C. Pulsed Interleaved Excitation. Biophys. J. 89, 3508–3522 (2005).

68. Schrimpf, W., Barth, A., Hendrix, J. & Lamb, D. C. PAM: A Framework for Integrated Analysis of Imaging, Single-Molecule, and Ensemble Fluorescence Data. Biophys. J. 114, 1518–1528 (2018).

69. Celetti, G. et al. The liquid state of FG-nucleoporins mimics permeability barrier properties of nuclear pore complexes. Journal of Cell Biology 219, e201907157 (2020).

70. Koshioka, M., Sasaki, K. & Masuhara, H. Time-Dependent Fluorescence Depolarization Analysis in Three-Dimensional Microspectroscopy. Appl. Spectrosc. 49, 224–228 (1995).

71. Szabelski, M. et al. Collisional Quenching of Erythrosine B as a Potential Reference Dye for Impulse Response Function Evaluation. Appl. Spectrosc. 63, 363–368 (2009).

72. Ranjit, S., Malacrida, L., Jameson, D. M. & Gratton, E. Fit-free analysis of fluorescence lifetime imaging data using the phasor approach. Nat. Protoc. 13, 1979–2004 (2018).

73. Anderson, J. A., Glaser, J. & Glotzer, S. C. HOOMD-blue: A Python package for high-performance molecular dynamics and hard particle Monte Carlo simulations. Comput. Mater. Sci. 173, 109363 (2020).

74. Dignon, G. L., Zheng, W., Kim, Y. C., Best, R. B. & Mittal, J. Sequence determinants of protein phase behavior from a coarse-grained model. PLoS Comput. Biol. 14, e1005941 (2018).

75. Togashi, Y. & Flechsig, H. Coarse-Grained Protein Dynamics Studies Using Elastic Network Models. Int. J. Mol. Sci. 19, 3899 (2018).

76. Varadi, M. et al. AlphaFold Protein Structure Database: massively expanding the structural coverage of protein-sequence space with high-accuracy models. Nucleic Acids Res. 50, D439–D444 (2022).

77. Varadi, M. et al. AlphaFold Protein Structure Database in 2024: providing structure coverage for over 214 million protein sequences. Nucleic Acids Res. 52, D368–D375 (2024).

78. Høie, M. H. et al. NetSurfP-3.0: accurate and fast prediction of protein structural features by protein language models and deep learning. Nucleic Acids Res. 50, W510–W515 (2022).

79. Barik, A. et al. DEPICTER: Intrinsic Disorder and Disorder Function Prediction Server. J. Mol. Biol. 432, 3379–3387 (2020).

80. Stukowski, A. Visualization and analysis of atomistic simulation data with OVITO–the Open Visualization Tool. Modelling Simul. Mater. Sci. Eng. 18, 015012 (2010).

