## supplemental table for "Differential conformational expansion of Nup98-HOXA9 oncoprotein in micro- and macrophases"

**Supplementary Table 1.** List of plasmids used in this study

| Name [vector_gene(s)] | Promoters | In Figure |
|---|---|---|
| pQE_14His::TEV::NHA9$^{\Delta GLEBS,\ A221TAG\text{-}S283Cys}$ | T5 | Fig. 1-4, Supplementary Fig. 2-6 |
| pQE_14His::TEV::NHA9$^{\Delta GLEBS,\ A221TAG\text{-}S312Cys}$ | T5 | Fig. 1-4, Supplementary Fig. 2-6 |
| pQE_14His::TEV::NHA9$^{\Delta GLEBS,\ A221TAG\text{-}S338Cys}$ | T5 | Fig. 1-4, Supplementary Fig. 2-6 |
| pQE_14His::TEV::NHA9$^{\Delta GLEBS,\ A221TAG\text{-}S362Cys}$ | T5 | Fig. 1-4, Supplementary Fig. 2-6 |
| pQE_14His::TEV::NHA9$^{\Delta GLEBS,\ A221TAG\text{-}S398Cys}$ | T5 | Fig. 1-4, Supplementary Fig. 2-6 |
| pQE_14His::TEV::NHA9$^{\Delta GLEBS}$ | T5 | Fig. 1-4, Supplementary Fig. 2-6 |
| pBI_Flag::NHA9::HA _Flag::NHA9::HA::P2AT2A::NHA9 | pBI_CMV | Supplementary Fig. 8 |
| pBI_Flag::NHA9$^{A221TAG}$::HA(boxB) _Flag::NHA9::HA::P2AT2A::NHA9 | pBI_CMV | Fig. 5, Supplementary Fig. 9 |
| pBI_Flag::NHA9$^{A221TAG\text{-}S283TAG}$::HA(boxB) _Flag::NHA9::HA::P2AT2A::NHA9 | pBI_CMV | Fig. 5, Supplementary Fig. 9 |
| pBI_Flag::NHA9$^{A221TAG\text{-}S312TAG}$::HA(boxB) _Flag::NHA9::HA::P2AT2A::NHA9 | pBI_CMV | Fig. 5, Supplementary Fig. 9 |
| pBI_Flag::NHA9$^{A221TAG\text{-}T353TAG}$::HA(boxB) _Flag::NHA9::HA::P2AT2A::NHA9 | pBI_CMV | Fig. 5, Supplementary Fig. 9 |
| pBI_Flag::NHA9$^{A221TAG\text{-}S362TAG}$::HA(boxB) _Flag::NHA9::HA::P2AT2A::NHA9 | pBI_CMV | Fig. 5, Supplementary Fig. 9 |
| pBI_Flag::NHA9$^{A221TAG\text{-}S98TAG}$::HA(boxB) _Flag::NHA9::HA::P2AT2A::NHA9 | pBI_CMV | Fig. 5, Supplementary Fig. 9 |
| pcDNA3.1_mEGFP::4×GGS::NHA9 | CMV | Fig. 5 |
| pcDNA3.1_TOM20$_{1\text{-}70}$::FUS$_{1\text{-}478}$::V5::Myc::4x$\lambda_{N22}$:: NES::PylRS$^{Y306A,Y384F}$_U6-tRNA$^{Pyl,CUA}$ | CMV, U6 | Fig. 5, Supplementary Fig. 9 |

**Supplementary Table 2.** Amino acid sequence of proteins used in this study

| Construct | Sequence |
|---|---|
| NHA9 (no GLEBS domain) | SGSMFNKSFGTPFGGGTGGFGTTSTFGQNTGFGTTSGGAFGTSAFGSSNNTGGLF<br>GNSQTKPGGLFGTSSFSQPATSTSTGFGFGTSTGTANTLFGTASTGTSLFSSQNNAF<br>AQNKPTGFGNFGTSTSSGGLFGTTNTTSNPFGSTSGSLFGPSSFTAAGPQNQVGAG<br>TTTGLFGSSPATSSATGLFSSSTTNSGFAYGQNKTAFGTSTTGFGTNPGGLFGQQN<br>QQTTSLFSKPFGQATTTQNTGFSFGNTSTIGQPSTNTMGLFGVTQASQPGGLFGTA<br>TNTSTGTAFGTGTGLFGQTNTGFGAVGSTLFGNNKLTTFGSSTTSAPSFGTTSGGL<br>FGFGTNTSGNSIFGSKPAPGTLGTGLGAGFGTALGAGQASLFGNNQPKIGGPLGT<br>GAFGAPGFNTTTATLGFGAPQAPVVDREKQPSEGAFSENNAENESGGDKPPIDPN<br>NPAANWLHARSTRKKRAPYTKHQTLELEKEFLFNMYLTRDRRYEVARLLNLTER<br>QVKIWFQNRRMKMKKINKDRAKDE |
| Flag::NHA9::HA::<br>P2AT2A::NHA9 | DYKDDDDKMFNKSFGTPFGGGTGGFGTTSTFGQNTGFGTTSGGAFGTSA<br>FGSSNNTGGLFGNSQTKPGGLFGTSSFSQPATSTSTGFGFGTSTGTANTLF<br>GTASTGTSLFSSQNNAFAQNKPTGFGNFGTSTSSGGLFGTTNTTSNPFGST<br>SGSLFGPSSFTAAPTGTTIKFNPPTGTDTMVKAGVSTNISTKHQCITAMKE<br>YESKSLEELRLEDYQANRKGPQNQVGAGTTTGLFGSSPATSSATGLFSSST<br>TNSGFAYGQNKTAFGTSTTGFGTNPGGLFGQQNQQTTSLFSKPFGQATTT<br>QNTGFSFGNTSTIGQPSTNTMGLFGVTQASQPGGLFGTATNTSTGTAFGT<br>GTGLFGQTNTGFGAVGSTLFGNNKLTTFGSSTTSAPSFGTTSGGLFGFGTN<br>TSGNSIFGSKPAPGTLGTGLGAGFGTALGAGQASLFGNNQPKIGGPLGTG<br>AFGAPGFNTTTATLGFGAPQAPVVDREKQPSEGAFSENNAENESGGDKPP<br>IDPNNPAANWLHARSTRKKRCPYTKHQTLELEKEFLFNMYLTRDRRYEV<br>ARLLNLTERQVKIWFQNRRMKMKKINKDRAKDEGGSGGSYPYDVPDYA<br>TGGSGATNFSLLKQAGDVEENPGPGSGEGRGSLLTCGDVEENPGPLQMF<br>NKSFGTPFGGGTGGFGTTSTFGQNTGFGTTSGGAFGTSAFGSSNNTGGLF<br>GNSQTKPGGLFGTSSFSQPATSTSTGFGFGTSTGTANTLFGTASTGTSLFSS<br>QNNAFAQNKPTGFGNFGTSTSSGGLFGTTNTTSNPFGSTSGSLFGPSSFTA<br>APTGTTIKFNPPTGTDTMVKAGVSTNISTKHQCITAMKEYESKSLEELRLE<br>DYQANRKGPQNQVGAGTTTGLFGSSPATSSATGLFSSSTTNSGFAYGQNK<br>TAFGTSTTGFGTNPGGLFGQQNQQTTSLFSKPFGQATTTQNTGFSFGNTST<br>IGQPSTNTMGLFGVTQASQPGGLFGTATNTSTGTAFGTGTGLFGQTNTGF<br>GAVGSTLFGNNKLTTFGSSTTSAPSFGTTSGGLFGFGTNTSGNSIFGSKPAP<br>GTLGTGLGAGFGTALGAGQASLFGNNQPKIGGPLGTGAFGAPGFNTTTAT<br>LGFGAPQAPVVDREKQPSEGAFSENNAENESGGDKPPIDPNNPAANWLH<br>ARSTRKKRCPYTKHQTLELEKEFLFNMYLTRDRRYEVARLLNLTERQVKI<br>WFQNRRMKMKKINKDRAKDE |
| mEGFP::4×GGS::<br>NHA9 | VSKGEELFTGVVPILVELDGDVNGHKFSVSGEGEGDATYGKLTLKFICTT<br>GKLPVPWPTLVTTLTYGVQCFSRYPDHMKQHDFFKSAMPEGYVQERTIF<br>FKDDGNYKTRAEVKFEGDTLVNRIELKGIDFKEDGNILGHKLEYNYNSHN<br>VYIMADKQKNGIKVNFKIRHNIEDGSVQLADHYQQNTPIGDGPVLLPDNH<br>YLSTQSKLSKDPNEKRDHMVLLEFVTAAGITLGMDELYKGGSGGSGGSG<br>GSMFNKSFGTPFGGGTGGFGTTSTFGQNTGFGTTSGGAFGTSAFGSSNNT<br>GGLFGNSQTKPGGLFGTSSFSQPATSTSTGFGFGTSTGTANTLFGTASTGT<br>SLFSSQNNAFAQNKPTGFGNFGTSTSSGGLFGTTNTTSNPFGSTSGSLFGPS<br>SFTAAPTGTTIKFNPPTGTDTMVKAGVSTNISTKHQCITAMKEYESKSLEE<br>LRLEDYQANRKGPQNQVGAGTTTGLFGSSPATSSATGLFSSSTTNSGFAY<br>GQNKTAFGTSTTGFGTNPGGLFGQQNQQTTSLFSKPFGQATTTQNTGFSF<br>GNTSTIGQPSTNTMGLFGVTQASQPGGLFGTATNTSTGTAFGTGTGLFGQ<br>TNTGFGAVGSTLFGNNKLTTFGSSTTSAPSFGTTSGGLFGFGTNTSGNSIFG |

| | |
|---|---|
| | SKPAPGTLGTGLGAGFGTALGAGQASLFGNNQPKIGGPLGTGAFGAPGFN<br>TTTATLGFGAPQAPVVDREKQPSEGAFSENNAENESGGDKPPIDPNNPAA<br>NWLHARSTRKKRCPYTKHQTLELEKEFLFNMYLTRDRRYEVARLLNLTE<br>RQVKIWFQNRRMKMKKINKDRAKDE |

Reference


1. Lakowicz, J. R. *Principles of Fluorescence Spectroscopy*. (Springer, 2006).
